# On the ancestry and evolution of the extinct dire wolf

**DOI:** 10.1101/2025.04.09.647074

**Authors:** Gregory L. Gedman, Kathleen Morrill Pirovich, Jonas Oppenheimer, Chaz Hyseni, Molly Cassatt-Johnstone, Nicolas Alexandre, William Troy, Chris Chao, Olivier Fedrigo, Savannah J. Hoyt, Patrick G.S. Grady, Sam Sacco, William Seligmann, Ayusman Dash, Mithil Chokshi, Laura Knecht, James B. Papizan, Tyler Miyawaki, Sven Bocklandt, James Kelher, Sara Ord, Audrey T. Lin, Brandon R. Peecook, Angela Perri, Mikkel-Holger S. Sinding, Greger Larson, Julie Meachen, Love Dalén, Bridgett vonHoldt, M Thomas P Gilbert, Christopher E. Mason, Rachel J. O’Neill, Elinor K. Karlsson, Brandi L. Cantarel, George R. R. Martin, George Church, Ben Lamm, Beth Shapiro

**Author notes:** Co-first authors (Gregory L. Gedman, Kathleen Morrill Pirovich).

## Abstract

Dire wolves (*Aenocyon dirus*) are extinct predators of Pleistocene North America. Although phenotypically similar to living wolves (*Canis lupus*), dire wolves have yet to be placed confidently in the canid family tree. We generated 3.4× and 12.8× paleogenomes from two well-preserved dire wolves dating to > 13,000 and > 72,000 years ago, and estimated consensus species trees for these and 10 canid species. Our results revealed that ∼2/3 of dire wolf ancestry is derived from a lineage sister to the clade comprising the gray wolf, coyote, and dhole, and the remaining ∼1/3 from a lineage near the base of Canini diversity. We identified 80 genes evolving under diversifying selection in dire wolves. Our results underscore the power of paleogenomes to resolve long-standing taxonomic questions and contribute to growing evidence of the role of post-speciation gene flow as an evolutionary force.

## INTRODUCTION

The dire wolf *(Aenocyon dirus*) first appeared in the North American fossil record during the late Middle Pleistocene ∼250,000 years before present (ybp) ^1^ and was the most prominent canid species in continental North America until its extinction ∼12,900 ybp (radiocarbon calibrated) ^2^. Their large size and morphological distinctiveness, as well as their abundance in fossil deposits at the Rancho La Brea Tar Pits (Los Angeles, California) has established dire wolves as iconic predators of the American Ice Age.

Dire wolves were morphologically similar to gray wolves (*Canis lupus*) with some exceptions. Dire wolves had larger temporalis muscles than gray wolves, capable of generating a greater bite force ^3^. They also had proportionately longer bacula, suggesting sexual competition ^4,5^. Dire wolves were comparable in size to the largest gray wolves – *C. l. occidentalis* in the northwestern United States ^6^ – and exhibited similar variation by latitude compared to gray wolves. Based on differences in size and geography, dire wolves are ascribed to two subspecies: the smaller ∼60 kg *A. d. guildayi* in the west, and the larger ∼68 kg *A. d. diru*s in the east ^7,8^.

Dire wolves are believed to have originated in North America ^9–11^, but their evolutionary history and taxonomic relationships to other canid species remain uncertain. A recent analysis of ∼0.01× to 0.23× coverage dire wolf genomes concluded that the dire wolf lineage first diverged from wolf-like canids, including gray wolves (*C. lupus*), coyotes (*C. latrans*), African golden wolves (*C. anthus*), golden jackals (*C. aureus*), Ethiopian wolves (*C. simensis*), dholes (*Cuon alpinus*), African wild dogs (*Lycaon pictus*), side-striped jackals (*Lupulella adustus*), and black-backed jackals (*L. mesomelas*), during the late Miocene, ∼6 million years ago (Mya) ^12^. This timing coincided with the first appearance in the Eurasian and African fossil records of the genus *Eucyon,* which is the lineage from which extant wolf-like canid species probably descended ^13^.

During the Pleistocene ice ages, shifting habitats and the ephemeral emergence of the Bering Land Bridge provided repeated opportunities for wolf-like lineages to disperse in and out of continental North America. Coyotes first appear in the North American fossil record 1-2 Mya ^14,15^, and gray wolves are present by the late Middle Pleistocene ^9,16^, although current North American gray wolf lineage entered North America from Eurasia around 25,000 years ago ^17–19^. Despite the geographic and temporal overlap between the ranges of dire wolves, coyotes, and gray wolves, no evidence of admixture between dire wolves and either lineage has been identified ^12^. Since admixture between closely related canid species is common ^20–22^, the absence of gene flow between dire wolves and sympatric canids suggests temporal, geographic, reproductive, or behavioral isolation.

Although Perri *et al.* ^12^ confirmed that dire wolves are a distinct evolutionary lineage, a high frequency of phylogenetic incongruence among estimated gene trees led to uncertainties about their early evolutionary history. About 50% of reported gene trees supported a topology in which dire wolves formed the earliest-diverging lineage of the wolf-like canid clade (Canina). Of the remaining gene trees, the side-striped and black-backed jackals were found to be an outgroup to a clade comprising dire wolves and other wolf-like canids ^12^. Divergence time estimates suggest the dire wolf lineage emerged after the divergence of the South American Cerdocyonina subtribe 5-4 Mya ^12,23^ and before the divergence of the common ancestors of African jackals (*Lupulella* spp.) 7.6-3.5 My and wolf-like canids 8.5-4.0 Mya (Canina subtribe). Incomplete lineage sorting ^24^ resulting from such rapid diversification could be one driver of this incongruence. In addition, while at least one admixture event between dire wolves and the common ancestor of dholes, coyotes, and wolves was detected, additional gene flow from unsampled populations, like the South American canids, could not be excluded.

Resolving the phylogenetic position of dire wolves and assessing evidence for gene flow is difficult given the millions of years of divergence between distinct canid lineages and technical challenges of analyzing ancient DNA. While low coverage paleogenomes can be useful for phylogenetic and demographic inference ^23,24^, ancient DNA is susceptible to DNA damage, deamination, and reference bias in mapping, in which divergent reads fail to map and similarity to the reference genome is artificially inflated ^25^. This bias results in an overestimation of sequence similarity between the ancient genome and the mapped reference, which can impact phylogenetic reconstruction and reconstructions of population history ^26,27^. This bias is particularly problematic when the reference genome is evolutionarily distant from the paleogenome ^28,29^, as is the case for the dire wolf.

Several factors contributed to the eventual extinction of dire wolves ^30,2,31^. The extinction of megafaunal prey species may have led to nutritional stress among dire wolves. The fossil record shows evidence of shrinking skull sizes during the Last Glacial Maximum ^32^, which can result from changes in diet and lifestyle. The loss of dire wolf prey and habitat from human-induced fires during the Late Pleistocene has been one proposed driver of these changes leading to their decline ^2^. Reduced population sizes may also have led to diversity loss and the accumulation of deleterious genetic variation due to increased rates of inbreeding. In support of this hypothesis, ossification disorders are observed in limb bones dating to several thousand years prior to dire wolf extinction ^31^.

To better understand the origin, evolution, and extinction of the dire wolf, we leveraged improved paleogenomics approaches and recovered additional ancient DNA from two well-preserved dire wolves ^12^: DireSP, an isolated incisor (Cincinnati Museum Center VP1737) recovered from Sheriden Pit, Ohio, estimated to be ∼13,000 years old based on calibrated radiocarbon dates from other bone material from the site ^33^. and DireGB, a complete isolated skull (Idaho Museum of Natural History, IMNH 48001/52) from Gigantobison Bay, a beach around the American Falls Reservoir, Idaho, dated to at least 72±14 thousand years ago based on the age of volcanic material from a nearby stratum ^34^. We generated paleogenomes for both specimens and performed phylogenetic, admixture, and selection analyses that provide new insights into the evolution and adaptation of the dire wolf to its unique niche.

## RESULTS

We generated paleogenomes with 3.4× depth of coverage of DireSP and 12.8× coverage of DireGB. After filtering, trimming, and merging overlapping reads, we retained 2.8 billion unique sequence reads for DireSP and 33.3 billion unique sequence reads for DireGB (**Table S1**). We mapped the reads using BWA *aln* against the Greenland gray wolf chromosome-level (including Y) genome assembly (GenBank GCA_905319855.2 aka “canLor”) ^35^ (**Table S2**). After mapping, 6% of the genome was covered >5× for DireSP and 82% for DireGB (**Fig. 1A-B**, **Table S3**). Previously, ancient DNA isolated from these same individuals yielded 0.2× (DireSP) and 0.3× (DireGB) coverage ^12^ (**Table S3**). We found patterns of DNA damage expected for single-stranded libraries (5’ C>T: DireGB = 30.2%; DireSP=26.6%, **Fig. S1**). Mapping to the gray wolf reference sex chromosomes versus autosomes yielded Rx ratios ^36,37^ indicating that both individuals were male (95% confidence intervals: DireSP=0.539-0.554, DireGB=0.538-0.554).

**Figure 1.**
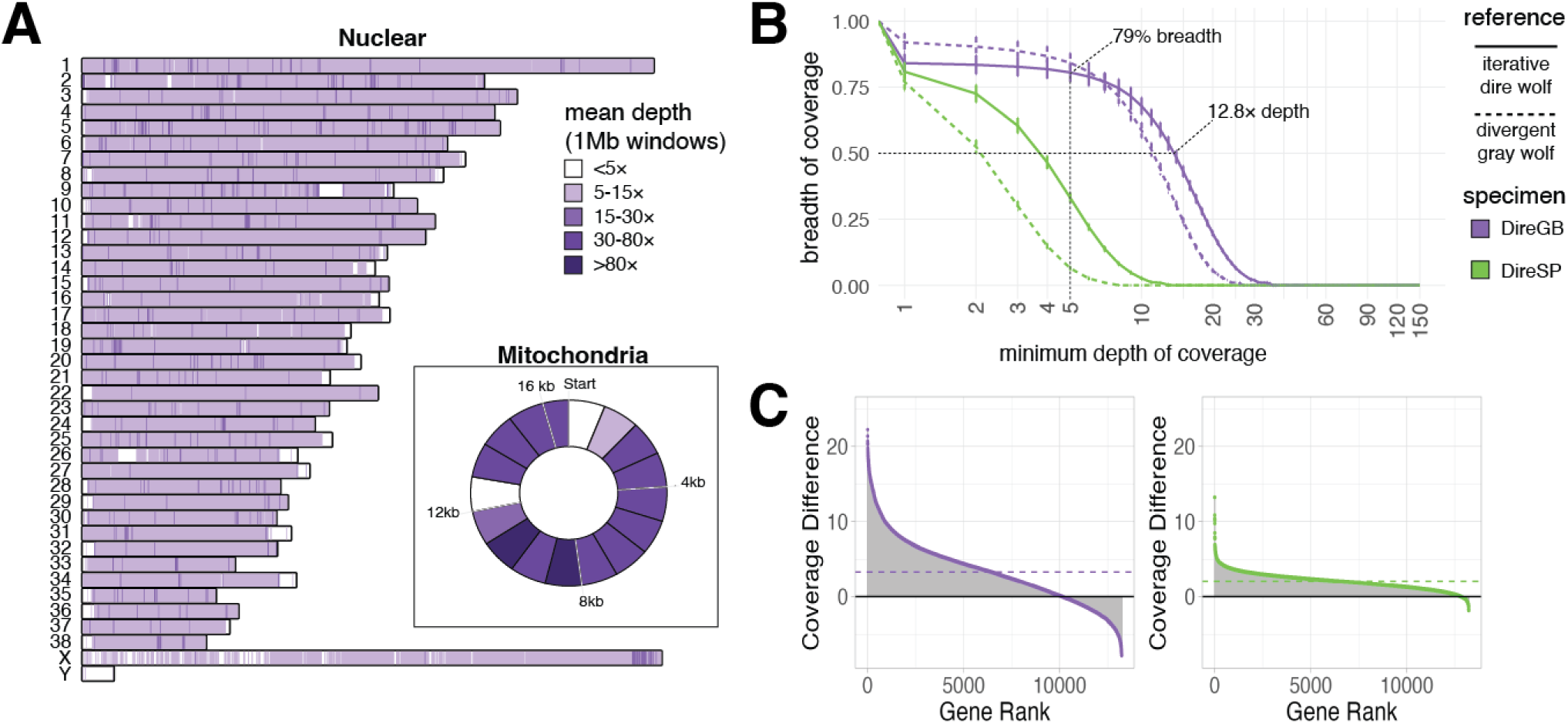
Iterative reconstruction of ancient dire wolf nuclear paleogenome improves whole genome and exome mappability. (**A**) Mean read depth of coverage by DireGB libraries (MQ>25) mapped over 1 Mb windows from the reference assembly of *A. dirus* reconstructed by iterative mapping and polishing. Inset displays mean read depth over the reconstructed full-length mitochondria sequence split by 1 Kb bins. (**B**) Cumulative coverage plot of DireGB (purple) and DireSP (green) libraries mapped to divergent gray wolf reference “canLor” (dotted line) and iterative dire wolf reference “aenDir” (solid line). Proportion of reference genome (i.e., breadth of coverage) at 5× and average depth for DireGB against the dire wolf reference are highlighted. (**C**) Average coverage difference over annotated orthologous genes (n=13,230) relative to dire wolf (positive) and gray wolf (negative). The majority of genes exhibited a boost in average coverage (dotted line) when using the reconstructed dire wolf paleogenome assembly in both DireGB (77% +3.3×) and DireSP (97% +2.1×).

We next used an iterative process to reconstruct a progressively less divergent nuclear paleogenome reference (“aenDir”) for mapping dire wolf ancient DNA sequencing libraries. After six iterations, we identified 14,569,091 changes (13,665,448 single nucleotide variants, 388,514 insertions, and 515,129 deletions) from the seed reference assembly (gray wolf). We then called a pseudo-haploid, variant-based consensus sequence that represented the highest depth alleles. In parallel, we used established methods ^28,38–40^ to reconstruct the mitochondrial paleogenome at high coverage (average depth = 158.4×). Merging both nuclear and mitochondrial reconstructed paleogenomes resulted in a final reference assembly with a hard-masked N content of 16.84% (**Table S4**).

Mapping ancient DNA libraries to the reconstructed paleogenome reference (aenDir) increased coverage both at the whole genome level and at the exome level (**Fig. 1A-B**). Compared with the initial mapping to the Greenland gray wolf genome reference, mapping to aenDir resulted in an increase in average coverage of 2.8× for DireGB and 1.5× for DireSP libraries. Following annotation of the reconstructed paleogenome (aenDir), we found that the majority of protein-coding genes (71%) were retained compared to the gray wolf reference (canLor), with most missing genes due to masking of low coverage regions (**Figs. S2,3**). The depth of coverage in exons improved by ∼2-fold for both DireSP and DireGB when mapped against the reconstructed dire wolf paleogenome (**Fig. 1C, Table S5**).

Mapping against the reconstructed paleogenome (aenDir) also reduced the impact of reference bias (**Fig. 2A**, **Table S6**). We found a significant decrease in allele sharing for the DireGB consensus called against the reconstructed dire wolf (aenDir) paleogenome compared to the gray wolf (canLor) genome, estimated using the *D*-statistic *D*(seed reference *Canis lupus*, non-seed reference *Canis lupus*; consensus *Aenocyon dirus*, outgroup *Vulpes vulpes*) (*D* for aenDir=-0.152, *Z*-score=-16.01; *D* for canLor=-0.521, *Z*-score=-76.51). In a complementary test, we assessed the identity-by-state (IBS) for DireSP reads mapped against the reconstructed aenDir paleogenome (**Fig. 2B**). We found that DireSP reads mapped to our reconstructed dire wolf genome exhibited higher IBS (i.e. greater sequence identity) compared to the gray wolf seed reference across all read lengths, especially for reads < 35 bp, suggesting the presence of residual reference bias, predominantly in shorter reads.

**Figure 2.**
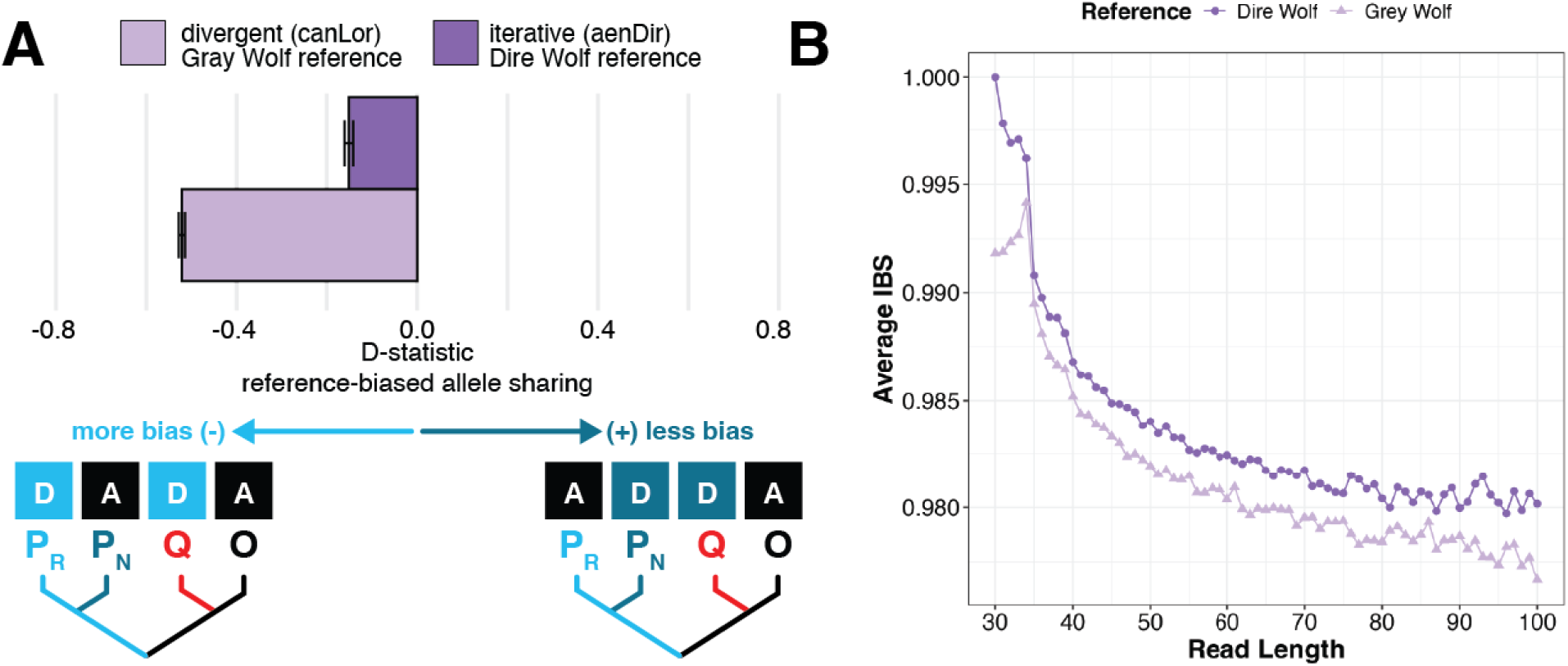
Iterative process to reconstruct the extinct dire wolf paleogenome sequence reduces reference bias. (**A**) Mapping to the reconstructed dire wolf paleogenome (aenDir) reduced reference bias 0.7 to 0.8 fold compared to mapping to the gray wolf reference genome (canLor). Residual reference bias was assessed using the ABBA/BABA test D statistic, which measures derived allele sharing of the DireGB consensus genome called against either the seed gray wolf reference genome (canLor) or reconstructed dire wolf paleogenome (aenDir) reference. Ancestral (A) and derived (D) alleles were localized by aligning either of the query (*A. dirus* in red, **Q**) consensus DireGB sequences (canLor- or aenDir-called), seed reference Greenland gray wolf (*C.lupus orion* in light blue, **P_R_**) and non-seed reference North American gray wolf (*C. lupus* in dark blue, **P_N_**) assemblies, and outgroup red fox assembly (*V. vulpes* in black, **O**) to a common reference genomic coordinates from a gray fox (*U.cinereoargenteus*) assembly. (**B**) Average identity-by-state (IBS) by read length for DireSP reads mapped to either the DireGB-based iteratively reconstructed paleogenome (dark purple) or divergent gray wolf (light purple) reference assemblies. IBS is defined as (matches - mismatch/aligned read length). Averages were calculated from a 500k read sample of the total alignment with an average of 5.7k (gray wolf) and 6.3k (dire wolf) reads per size bin.

Dire wolf paleogenomes were unexpectedly more similar to extant Canina genomes than anticipated given a 5.8 Mya divergence time ^12^, a mutation rate calibrated for *Canis* species ^41^, and a wide range of generation times ^42^ for priors (**Fig. S4**). We computed sequence divergence on fixed autosomal transversions at sites covered at 5× relative to the total non-N sequence length of the reconstructed dire wolf paleogenome reference (2,310,286,013 bp). We found the dire wolf paleogenomes less diverged (on average, 0.09% divergent) than expected based on prior parameters (0.6% to 0.9% sequence divergence), and estimated sequence similarities of 99.91% to jackals and 99.94% to non-seed reference gray wolves at either end of wolf-like canid diversity (**Fig. S4A**). As residual reference bias in mapping could artificially inflate sequence similarity, we also assessed the distribution of unique 32-mers from the contamination-filtered sequence reads using *mash* for metagenome distance estimation ^43^. We found that dire wolves exhibited equally low 32-mer diversity compared to gray wolf and jackal species (∼0.01), with greater distance to red and gray foxes (0.02 to 0.03) (**Fig. S4B, Table S7**).

Our estimated mitochondrial and nuclear phylogenies were similar to those published previously (**Figs. 3, S5-9, Tables S8-10**). The mitochondrial phylogeny showed the same mitonuclear discordance that is known to characterize the canid lineage ^22^. Our nuclear phylogeny, which was based on 19.5Mb of non-coding dire wolf sequence at least 1Mb away from protein-coding genes and <25% missingness on average (**Figs. S6,7**), placed dire wolves into a monophyletic clade that diverges prior to the split between the black-backed jackal and wolf-like canids (**Figs. 3, S8**). However, this placement was only supported by 44% of input loci, while 39% of loci supported a closer relationship between dire wolves and non-Lupulellan wolf-like canids (**Fig. 3**). Divergence time estimates (**Fig. S9, Table S10**) suggested that the dire wolf diverged from other canids ∼4.5 million years ago (95% CI=5.84-2.89 Mya) and the jackals diverged ∼4.08 million years ago (95% CI=5.34-2.58 Mya).

**Figure 3.**
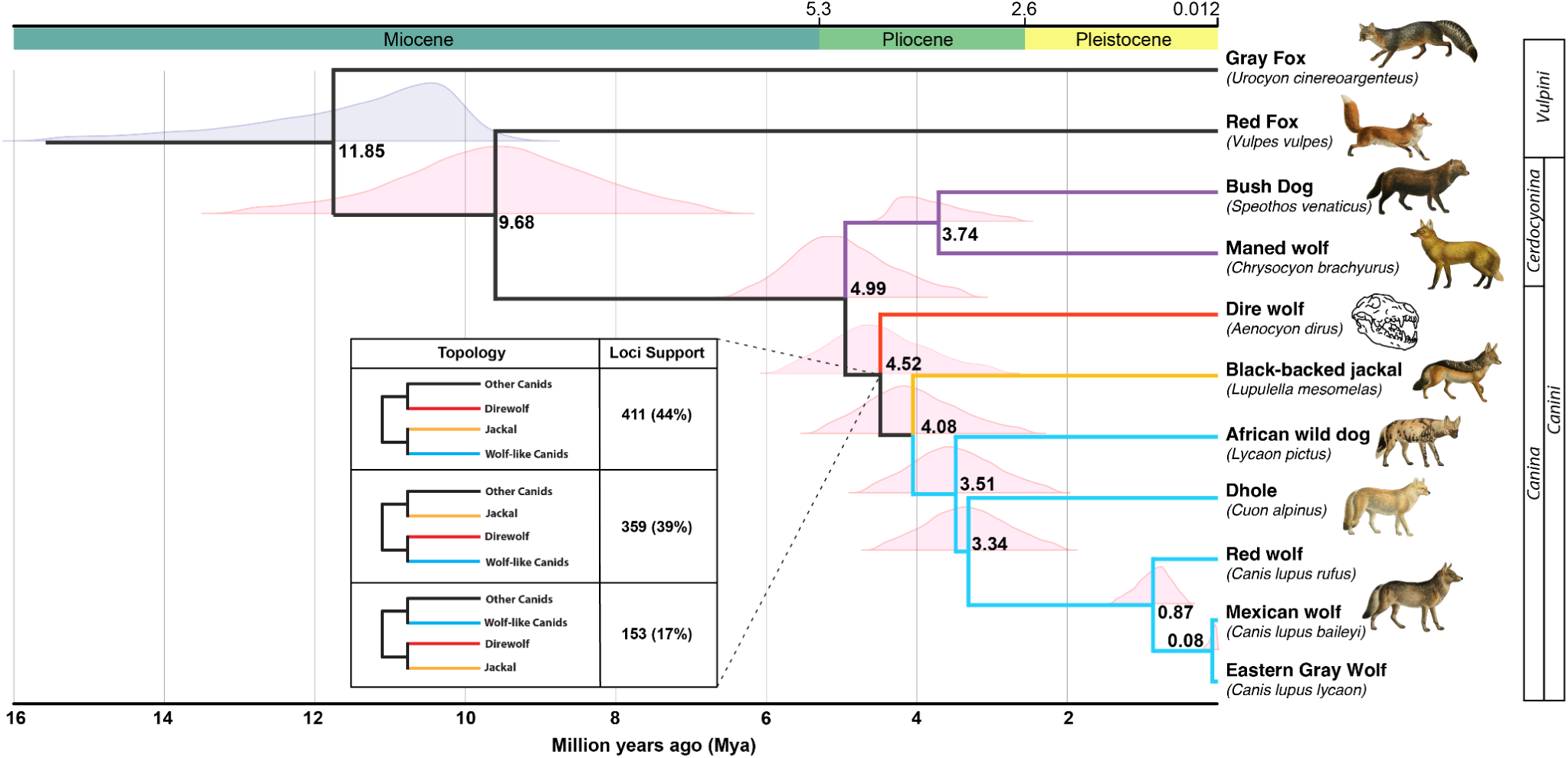
The dire wolf lineage is distinct from jackals and other wolf-like canids. Time-calibrated nuclear phylogenetic relationship of extinct dire wolves to extant canids using MCMCtree from 25 kb loci trees (n=924) using IQTree and species tree consensus using ASTRAL. Dire wolves (red) diverge prior to the split of black-backed jackals (yellow) and wolf-like canids (blue). Root ages are mean divergence time estimates in millions of years (Mya). Node distributions visualize confidence in placement. Node distributions with fossil calibrations are shown in purple. Inset details the alternative topologies for the dire wolf node, along with the proportion of loci supporting each alternate.

The discordance among estimated gene tree topologies appears to arise from admixed ancestry in dire wolves (**Fig. 4**). Dire wolves shared an excess of derived alleles with gray wolves relative to other canids, including dholes (11.9 ≤ *Z* ≤ 15.3), African wild dogs (61.9 ≤ *Z* ≤ 65.4), African jackals (92.3 ≤ *Z* ≤ 99.6), and South American canids (173.4 ≤ *Z* ≤ 183.7) (**Fig. 4A**). We did not detect similar affinity between dire wolves and gray wolves when comparing to wolf-like canids that are more closely related gray wolves, including coyotes and red wolves (*|Z|* ≤ 2.4) (**Fig. 4A**). We also observed that, relative to dire wolves, African jackals (66.0 ≤ *Z* ≤ 72.4) and other wolf–like canids (*Z* ≥ 92.7) share more derived alleles with gray wolves (**Fig. 4B**). Together, these patterns suggest that dire wolves had a mixture of ancestries from two distinct lineages, with one diverging prior to African jackals and another stemming before gray wolves and coyotes. Allele sharing between dire wolves and South American canids relative to extant wolf-like canids (-7.2 ≤ *Z* ≤ -2.3) is consistent with the deeper ancestry component falling within Cerdocyonina diversity (**Fig. 4B**). Both dire wolves formed a clade relative to all individual coyotes and gray wolves (**Fig. S10**), implying that there was limited gene flow between dire wolves and specific *Canis* lineages following their divergence.

**Figure 4.**
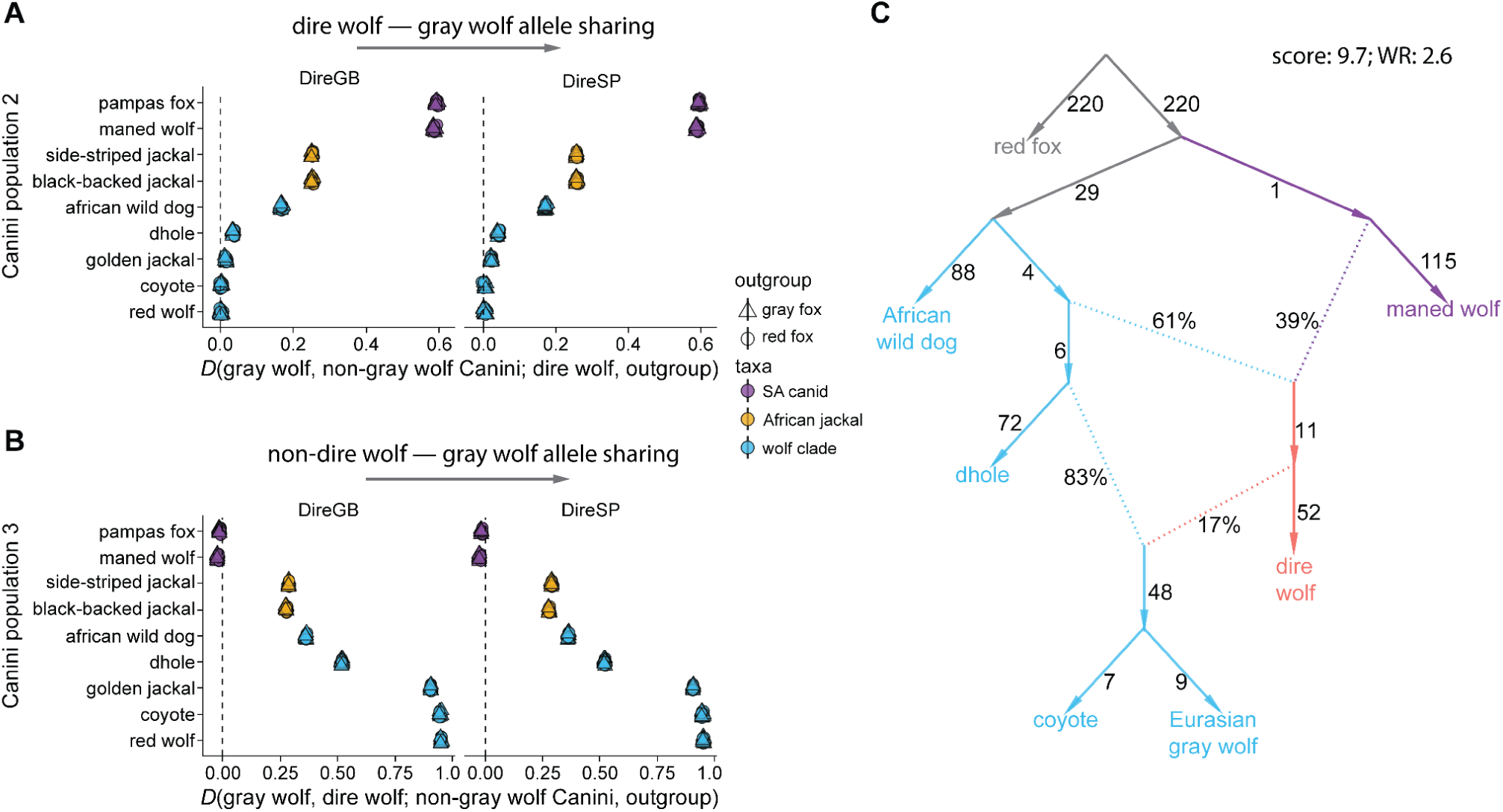
Investigating gene flow between dire wolves and other canid lineages. (**A**) D-statistics of the form *D*(gray wolf, other Canini group; dire wolf, red fox), investigating allele sharing between dire wolves and gray wolves, relative to other species and subspecies within Canini. (**B**) *D*-statistics of the form *D*(gray wolf, dire wolf; other Canini group, red fox), investigating allele sharing between dire wolves and other populations within Canini, relative to gray wolves. Bars depict 3 standard errors in (**A**) and (**B**). Each point is a comparison with a different individual gray wolf, with points colored by the Canini clade being of the non-gray wolf population in the comparison. (**C**) Best-fitting admixture graph with two admixture events modeling the relationship between dire wolves (n=2); wolf-like canids: Eurasian wolves (n=2); coyotes, (n=4), dholes (n=2), African wild dogs (n=3); and maned wolf (n=2); with red fox (n=2) as the outgroup. Additional graph fits are summarized in Table S11.

Admixture graph models of canid population history identified both wolf-like canid and deep ancestry components in the dire wolf lineage (**Fig. 4C**). We applied a heuristic algorithm ^44^ to find optimal admixture graphs modeling the relationships between seven canids (dire wolves, Eurasian gray wolves, coyotes, dholes, African wild dogs, maned wolves, using red foxes as an outgroup). No model permitting zero or one admixture event was supported by the data; however, the best-fitting model featuring two admixture events had statistical support (worst residual of *Z*=2.6) (**Fig. 4C**). This model found that dire wolves received a large portion of their ancestry (∼39%) from a lineage that groups with South American canids, with the remaining ancestry (∼61%) attributed to a sister lineage to dholes, coyotes, and gray wolves. This wolf-like ancestor may have been the source of the phylogenetically-discordant dire wolf mitochondrial genome, which groups within wolf-like canid diversity. When analyzed separately, both dire wolf individuals produced similar optimal graphs (**Fig. S13**), suggesting that admixture was a species-wide feature of dire wolf population history. Graphs that include both an additional outgroup (gray foxes) and additional South American canid species confirmed that the divergent portion of dire wolf ancestry falls near the base of Canini diversity (**Fig. S13**).

The phylogenetic placement of the wolf-like component in dire wolves is remarkably similar to a proposed unsampled canid lineage that contributed ancestry to the common ancestor of coyotes and wolves ^45^. The best-fitting model similarly features gene flow from dire wolves into wolves and coyotes, which could be the source of this unsampled ancestry. However, scenarios in which dholes, rather than coyotes and wolves, are admixed also provide similar fits to the data (**Table S11**) and we observed that relationships among extant wolf-like canids can be modeled without requiring this unsampled admixture (**Fig. S12**). As suggested by evidence of excess derived allele sharing between dire wolves and South American canids, a similar model which instead places the minor ancestry component of dire wolves outside extant Canini diversity instead of on the South American canid branch provides a worse fit for the data (worst residual of *Z*=5.25). Models of canid population history that include additional extant canid lineages highlight other known and cryptic admixture events and identify the gene flow event leading to dire wolves (**Fig. S14**). We found mixed support for introgression from the dhole lineage into the African wild dog lineage, which could reflect cryptic admixture ^45,46^, and consistent support for other previously detected admixture events ^20,45,47,48^. These findings reinforced that this approach is sensitive enough to detect hybridization across diverse canids (**Fig. S18**), and does not appear to be impacted by choice of reference genome (**Fig. S13**).

The ubiquity of gene flow among canids, especially among diverged wolf-like canid species ^45–49^, poses challenges for inferring the precise placement of admixture events among these taxa. Our goal was to infer the simplest models which described the data, both for the minimal (**Fig. 4C**) and expanded sets of canids (**Fig. S14**), which each provided evidence for the dual ancestry of dire wolves. However, given extensive hybridization among canids, the true population history may be more complex. More complicated models, such as those featuring three admixture events in the minimal canid species set, also largely suggest hybridization in dire wolves, though a much wider range of graphs fit the data, including many which have other lineages, rather than dire wolves, as admixed (**Table S11**). Challenges arising from mapping sequence data diverged taxa also complicate inferences of canid population history. We observed several biases across alignment and genotype calling strategies (**Fig. S10, S19**). The Consensify-style pseudohaploid genotype calls using pangenome alignments presented here appeared to minimize these biases (**Fig. S13, S19)**, though do require greater agreement between reads, which has the potential to amplify biases arising in mapping.

Comparative sequence analysis detected molecular signatures for diversifying and adaptive evolution in dire wolf protein-coding genes (**Fig. 5**). We used the HyPhy framework ^50–52^ to identify genes with signatures of positive selection among dire wolves and 10 other canid taxa, including episodic diversifying selection in dire wolves contrasted to other canids. We identified 49 genes under significant (BUSTED *p* < 0.05) episodic positive selection and another 31 genes under non-episodic positive selection in the dire wolf branch (free-ratio dN/dS > 2.0) (**Fig. 5, Table S12**). None of these genes exhibited pervasive positive selection over the canid tree (one-ratio dN/dS < 2.0), although dire wolves may share positive selection at a site level with other branches. Rarely, we found that the dire wolf codon matched a codon found in another branch. For example, the 505th codon of *NCAPG* (MEME *p*= 3.31e-06) had two branches under positive selection: the root codon encodes serine (TCT) but both the dire wolf and *Canis* branches, and not other wolf-like canids, encode asparagine (AAT). Likewise, the 794th codon of *WNK4* (MEME *p*= 0.00513), encoded asparagine (AAC) in both dire wolf and *Lycaon* branches rather than the root alanine (GCC). We did not find extensive functional enrichment for gene sets across the 80 genes under diversifying positive selection in the dire wolf branch. Functional domain analysis ^53^ and protein language modeling ^54^ identified 42 dire wolf codons in 22 genes under episodic diversifying selection that intersect protein functional domains which could be impacted by amino acid substitution (**Table S13**).

**Figure 5.**
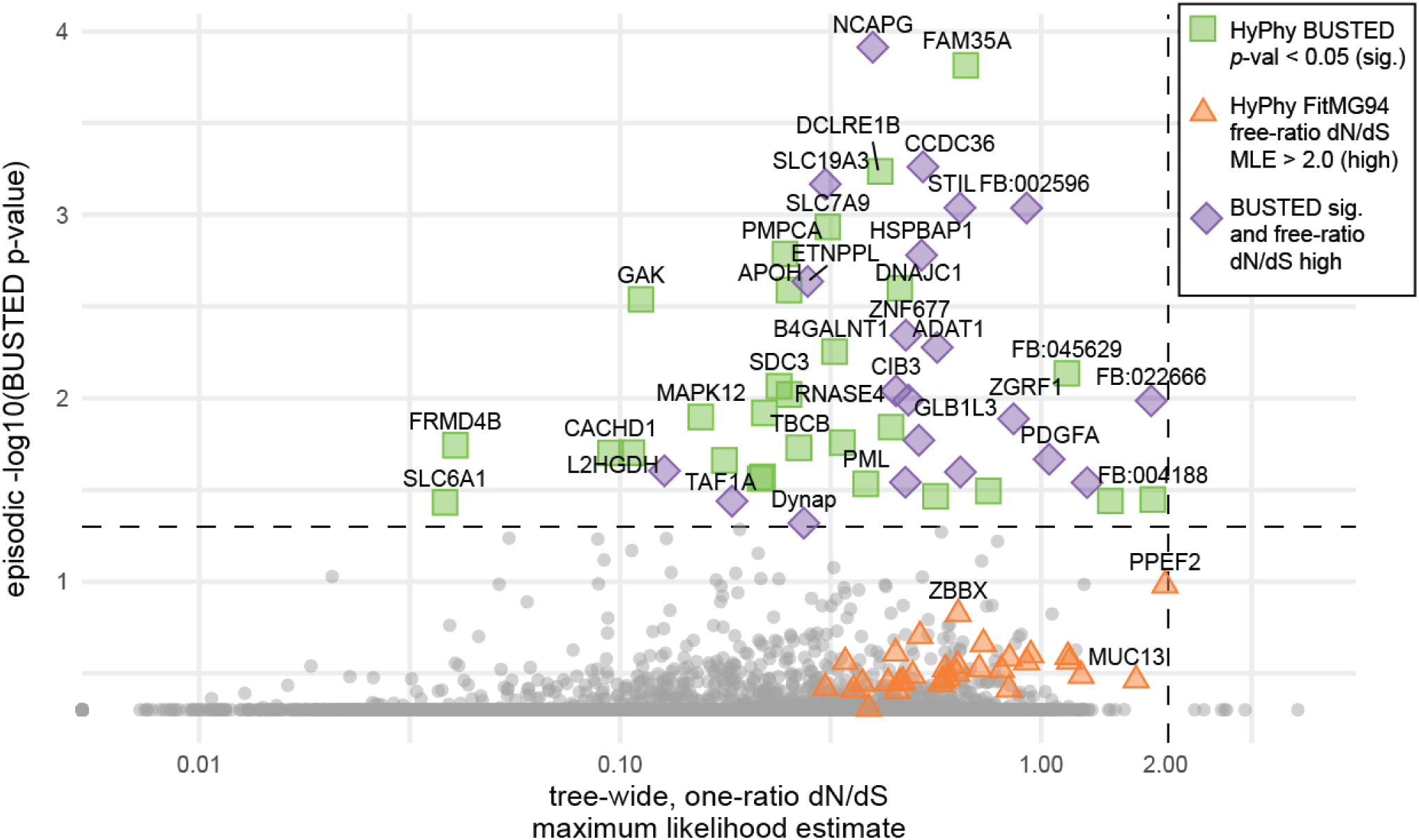
Comparative analysis of nuclear protein-coding gene sequences supported by dire wolf ancient DNA and 10 other canid genomes inform pathways under episodic or pervasive diversifying positive selection that may shape species adaptations and phenotypes. The -log10 HyPhy BUSTED *p*-values for episodic positive selection in the dire wolf branch plotted (y-axis) over the canid tree-wide, one-ratio dN/dS ratio maximum likelihood estimates (x-axis) for 5,984 genes where high one-ratio dN/dS implies pervasive positive selection across the canid lineage. We find 31 genes exhibiting branch-specific, free-ratio dN/dS > 2 (orange triangles), 28 genes significant for branch-specific, episodic selection in BUSTED test *p* < 0.05 (green squares), or 21 genes satisfying both conditions (purple diamonds). Annotated genes without gene symbols of orthologs identified are labeled with “FB:”.

## DISCUSSION

### Dire wolf evolutionary history

The Late Pleistocene dire wolf is descended from a canid lineage that emerged following admixture between two lineages: a lineage along the branch that gave rise to the South American canids (Cerdocyonina subtribe) after they diverged from African jackals and other wolf-like canids (Canina subtribe), and a more recently derived wolf-like canid lineage that diverged after the emergence of the African wild dog (*Lycaon*) lineage (**Figs. 3**, **4C**). This admixed ancestry explains the discordance in nuclear gene phylogenies observed both here and previously reported in Perri et al. (2021).

Given fossil evidence that dire wolves were only present in North America, the ancestral components of the dire wolf lineage might be attributed to other canids in the North American fossil record. The oldest wolf-like fossils in the North American fossil record are from the tribe Canini ^10^, including the extinct genus *Eucyon* (10.3-3.6 Mya), the proposed progenitor of *Canis* and *Lupulella*, and the earliest ancestors of the Cerdocyonina subtribe, which dispersed to South America ^55–57^. It would be plausible, given an estimated divergence timing straddling Late Miocene and Pliocene, that the major ancestry component of the dire wolf is from an early North American *Eucyon* species, but our admixture graph modeling supports instead a major ancestor diverging after the *Lycaon* lineage (African wild dogs), which would be post *Eucyon* expansion outside of North America*. Eucyon* expanded into Eurasia and Africa during the Late Miocene, where it gave rise to *Lupulella*, the African jackals ^57–61^, as well as *Canis* and the related genera *Lycaon* and *Cuon* ^46^. From the late Miocene through the Pleistocene, various *Canis* species expanded back into North America, supplanting or admixing with local populations ^13^. *Canis lepophagus* expanded into the Americas during the late Miocene to early Pleistocene ^10,62^, for example, followed by *C. armbrusteri* and *C. edwardii* in the Pliocene and Pleistocene. The wolf-like lineage that contributes majority ancestry to dire wolves could be one of these earlier expanding lineages.

We note that the deep divergence (∼10 Mya) among the canid lineages examined here makes it challenging to examine gene flow across these species while accounting for reference bias, particularly with highly fragmented ancient DNA. While our iterative mapping approach mitigated reference bias in our alignment of dire wolf sequences, it is possible that our more fragmented dire wolf reference reduced mapping performance and amplified other biases in the alignment of short reads from non-dire wolf canids. Thus, we complemented analyses of canid population history with a pangenome-mapping approach which mitigated these biases. Additionally, the prevalence of interspecific gene flow among canids makes it unlikely that any particular model will fully capture the underlying population history relating these species ^47^. However, our conclusion regarding the dual ancestry of dire wolves is robustly supported across different models and approaches.

Our results underscore the role of post-divergence gene flow as an evolutionary force shaping species across the tree of life, and the potential for lineages to be forged through hybrid speciation – as uncovered in the Columbian mammoth (*Mammuthus columbi*), the same megafauna that roamed alongside dire wolves ^63,64^. For example, excess basal ancestry in wolves and coyotes relative to other wolf-like canids was previously attributed to gene flow with an unknown wolf-like canid lineage that emerged following the divergence of African wild dogs ^65^, raising the possibility that the same unknown lineage interbred with both groups. Indeed, our results suggest that the wolf-like lineage found within dire wolves also contributed ancestry to wolves and coyotes, possibly mediated through gene flow with dire wolves. However, we infer that gene flow between diverged canid lineages was common, and that the relationships within the wolf-like canids can be modeled without invoking this unknown lineage. Our admixture graph analyses do, however, uncover previously described episodes of gene flow between canids, resulting in admixed ancestries for species including eastern coyotes ^48^, red wolves ^47^, golden wolves ^20,45^, and dholes ^46^.

### Evolutionary adaptations of dire wolves

We identified several genes in the dire wolf genome that may have contributed to one of their primary evolutionary features: their large builds. *NCAPG,* a gene associated with body size, was the most significant gene for episodic diversifying selection in the dire wolf branch. Yet, dire wolves and gray wolves exhibit unusually low rates of synonymous substitutions in *NCAPG* (both dS= 4e-10) – magnitudes lower than other canids (all dS > 0.001) – and together share a serine-to-asparagine substitution under site-specific positive selection (MEME *p*= 3.31e-06). This degree of sequence constraint could suggest strong purifying selection ^66–69^, perhaps upon non-coding regulatory elements influencing expression of adjacent gene *LCORL*, just 200 kilobases away. *LCORL* is an allosteric regulator of the polycomb repressor complex (PRC2) ^70^, and haplotypes spanning *NCAPG* and *LCORL* have been strongly associated with growth traits in domestic dogs^71^, goats^72^, sheep^73^, horses^74^, and cattle^75,76^. Expression levels of *NCAPG* and *LCORL* have inverse relationships on growth: positively correlated for *NCAPG* levels ^77^ and negatively correlated for *LCORL* levels ^74^. We hypothesize that in both dire wolves and gray wolves, purifying selection has maintained ancestral regulatory elements at the *NCAPG* locus, while in other wolf-like canids of smaller body sizes, diversifying selection altered co-regulation of these genes.

We also found several genes under episodic diversifying selection in dire wolves annotated as affecting germ cell development, gamete function, and reproductive tissue structure, which may contribute to the reproductive biology of dire wolves distinct from wolf-like canids ^4^. The genes *B4GALNT1*, *ROS1*, *TP73*, *AKAP3*, *NUTM1*, and *ABCG5* are all associated with “male infertility” in the Mammalian Phenotype Ontology (MP:0001925). While most of these fertility-related genes have not been implicated in reproductive structures, *ROS1* regulates the differentiation of male epididymis ^78^. *PDGFA*, another gene under positive selection in dire wolves, has been shown as integral to epididymal morphology in mice ^79^. None overlap the few genes associated with mammalian baculum size, however ^80^, meaning that genes shaping divergent reproductive biology in dire wolves are still a mystery.

### An iterative approach to nuclear genome inference

We showed that iterative mapping combined with conservative consensus calling can help mitigate the impacts of reference bias and DNA damage in paleogenomics. To generate the dire wolf paleogenome, we mapped reads to the Greenland gray wolf as a seed reference, called consensus from aligned dire wolf reads, then re-mapped reads to this new consensus. We then repeated this process until no additional reads were added. Our intent was to estimate a conservative assembly, and thus we chose to hard-mask bases (i.e., Ns) not supported by >5× coverage, resulting in a final reference assembly comprising only the most abundant alleles from mapped dire wolf reads. Methods like EAGER use a similar conservative consensus calling approach when mapping to the nuclear genome, but employ only a single round of mapping and thus do not capture the maximum number of reads ^81^. Iterative assembly tools are used in ancient DNA research, but mostly for mitochondrial genome assembly (e.g., MIA ^82^) as they do not scale to whole nuclear genomes ^83^. Iterative mapping was used to reconstruct the Denisovan genome, where Meyer et al. (2012) ^38^ performed two rounds of iterative mapping and then filled in low coverage regions with human sequence, which is a source of reference bias. Given the improvements that we observed with multiple rounds of iterative mapping, we recommend a similar approach, in particular when no close reference genome is available.

One concern with an iterative mapping approach is that the process could actually exacerbate reference bias rather than reduce it. If the seed reference is too divergent, for example, only the most similar ancient reads will map, potentially masking informative sites and compounding bias. We approximated reference bias using *D*-statistics ^38,84^ by assessing derived allele sharing with distinct gray wolf subspecies, and found no evidence of compounding bias. In fact, we observed a reduction in bias towards the seed reference Greenland gray wolf genome. Since shorter reads have been shown to be more impacted by reference bias ^40,85^, we performed a secondary assessment of identity by state (IBS) by read length from an unrelated dire wolf individual (DireSP). While residual reference bias remains, particularly for shorter reads, we observed higher IBS on average for all DireSP reads regardless of length, suggesting we successfully generated a less divergent reference genome that facilitated better mapping of ancient reads.

While our approach estimated a paleogenome that was more similar to the dire wolf relative to the seed reference, steps could be taken to improve these genomes further. For example, while our approach can identify and incorporate SNPs and small insertions or deletions into the final reference, it cannot account for larger structural variants specific to the dire wolf. The reconstruction of large structural variants is unresolved for short read data sets that are typical of paleogenomes ^86^ and, given our average read length of 69 bp across all samples, is not achievable for the dire wolf. Nevertheless, by iteratively estimating a paleogenome reference, we could recover more reads that would have otherwise missed mapping to a divergent reference genome without those small sequence differences.

An alternative – or, perhaps, complementary – approach to mitigating reference bias is to construct a pangenome from multiple divergent species for initial mapping ^87,88^, which allows for capture of highly divergent reads and, potentially, larger structural variants. The benefits of a pangenome approach were demonstrated by Martiniano and colleagues (2020) ^89^, who saw a reduction of reference bias in mapping simulated ancient reads to the 1000 Genomes Project variational graph. By mapping real reads from ancient humans to this pangenome, they could capture sequence variation missed by initial mapping to a single reference without a loss of mapping sensitivity. However, divergent dire wolf haplotypes not represented by the available paths in the pangenome will still be affected by reference bias. Nevertheless, cost-efficient optimizations that enable a combination of iterative mapping and polishing approach will provide additional improvements to estimating increasingly accurate paleogenomes from short read data sets.

These data provide a comprehensive overview of dire wolf genomics, evolution, and adaptations, offering insights into this iconic species. The reconstructed reference paleogenome provides a resource for the broader research community and a roadmap for future studies aiming to leverage high-coverage ancient DNA. The genes identified under selection in dire wolves offer insights into possible molecular mechanisms for this species’ role as a predator of extinct megafauna. This work furthers our understanding of the dire wolf and their place in the broader patterns of canid evolution.

## STAR METHODS

### Published ancient DNA sequencing data

We downloaded ancient DNA sequencing reads from dire wolf individuals previously sampled, including the Sheriden Pit individual (DireSP, BioSample accession SAMEA5559854) and the Gigantobison Bay individual (DireGB, BioSample accession SAMEA5559855) and three other individuals from American Falls Reservoir, Idaho (DireAFR, BioSample accession SAMEA5559852), Natural Trap Cave, Wyoming (DireNTC, BioSample accession SAMEA5559856), and Guy Wilson Cave, Tennessee (DireGWC, BioSample accession SAMEA5559853) publicly released under BioProject accession PRJEB31639 (**Table S1**).

### Provenance and sampling of dire wolf individuals

We obtained permission to collect new samples from two dire wolf individuals that previously yielded authentic dire wolf mitochondrial and nuclear DNA ^12^. The first dire wolf, referred to as “DireSP” (Cincinnati Museum Center; VP1737; ACAD1735; DireSP), is from Sheriden Pit site in Indian Trail Caverns, Ohio, and an incisor tooth previously sampled at the root (BioSample accession SAMEA5559854). We sampled an additional incisor from this individual for the present study. The second dire wolf, referred to as “DireGB” (Idaho Museum of Natural History; IMNH 48001/52; ACAD18742; DireGB) is from Gigantobison Bay site at American Falls Reservoir, Idaho, and presents a complete cranium with intact teeth. The left petrosal was previously sampled (BioSample accession SAMEA5559855) and the right petrous was sampled for this study. The Sheriden Pit individual could have been deposited before 13,000 years ago, based on calibrated radiocarbon dates from other bone material derived from the same layer ^33^, but has not been radiometrically dated, and the Gigantobison Bay individual is probably older than 72,000 years, based on the age of volcanic material from a nearby stratum ^34^.

### Sample preparation and ancient DNA extraction

For the Sheriden Pit individual (BioSample accession SAMN46779696, we extracted ancient DNA from an incisor (Cincinnati Museum Center VP1737). We used a Dremel with drill attachment to obtain bone powder from the interior of the root. We rinsed the obtained bone powder in a 0.5% bleach solution for fifteen minutes followed by three water washes before proceeding with extraction. For the Gigantobison Bay individual (BioSample accession SAMN46779697), we extracted ancient DNA from the incus bone recovered from the right petrous of a complete skull (IMNH 48001/52). Ossicles are an optimal source of DNA preservation ^90,91^. We rinsed the whole incus in a 0.5% bleach solution, followed by three water washes. We extracted DNA for both individuals using the Buffer D option, a protocol optimized for recovering short, degraded molecules ^92^. For the Sheriden Pit sample, we extracted DNA from two 5 mg aliquots of bone powder following bleach pre-treatment. From these two DNA extracts we generated three single stranded libraries each ^93^. Following the initial round of extractions we observed a non-dimineralized bone powder pellet in one of the two lysate tubes and performed a re-digestion of this bone powder following Rohland et al. (2018) ^92^. From the DNA isolated after this re-digestion we generated two more single-stranded libraries ^93^. For the Gigantobison Bay individual, we extracted DNA from the whole incus, which was completely dissolved following overnight lysis. We then prepared single-stranded libraries from the extracts as per Kapp et al. (2021) ^93^.

We generated a single extract from the Gigantobison Bay ossicle, from which we made six libraries. All pre-PCR work, including sample preparation, DNA extraction, library construction, and quality control sequencing, was carried out in a dedicated ancient DNA lab at the University of California, Santa Cruz.

### Massively parallel paired-end short DNA read sequencing

We prepared a total of nine Illumina libraries from the sample of the Sheriden Pit dire wolf (DireSP) and six Illumina libraries from the sample of the Gigantobison Bay dire wolf (DireGB). We sequenced all libraries on a Nova-seq 6000 using version 1.5 chemistry, loading 0.3nM final library concentration. For the DireSP sample, we obtained 1,320gb (R1+R2) of raw sequencing data or 13.2 billion total paired end reads (R1+R2). For the DireGB sample, we obtained 7,814gb (R1+R2) or 78 billion total paired end reads (R1+R2). We report the total number of raw sequencing reads generated per run, library, and individual in **Table S1**. For subsequent steps of parallelized processing, we split sequencing reads into batches of 50,000,000 reads.

### Sequence read pre-processing and quality control

We trimmed raw sequencing reads using *fastp* ^94^ (*i*) to remove adapter sequence, low quality ends of reads, and reads less than 35 nucleotides, (*ii*) merge and correct overlapping reads with a minimum of 15 nucleotide overlap, and no more than three differences, and (*iii*) trimmed homopolymer G and X sequences at the 3’ end (-q 25 -l 35 -m --overlap_len_require 15 --overlap_diff_limit 3 -c -g - x). We classified sequence contaminants using *Kraken 2* ^95^ and a confidence of 0.8 against the pre-compiled 8 GB database MiniKraken DB_8GB constructed from complete bacterial, archaeal, and viral genomes in RefSeq (2017-10-18). We used combine_kreports.py from *Kraken Tools* ^96^ to merge all metagenomic classification reports generated for split read batches. We report statistics on sequencing reads before and after processing for both published and novel ancient DNA sequencing libraries in **Table S1**.

### Reference genome assemblies and annotations

We selected the Greenland gray wolf (*Canis lupus*) chromosome-level (including Y) genome assembly (GenBank GCA_905319855.2) ^35^ as our primary single-reference genome assembly of a modern canid species, and utilized additional canid genome assemblies ^97–114^ throughout this study as referenced in **Table S2**. We surmised that the Greenland gray wolf would serve as a suitable scaffold reference due to the geographic isolation of Greenland wolves relative to other gray wolves ^115^, the fully assembled chromosomes including Y and high contig N50 relative to other canid genome assemblies (**Table S2**), and to previously inferred phylogenies that suggest it is no more distantly related to dire wolves than any other extant members of the Canina subtribe.

### Read mapping to single reference genomes

We utilized two primary mapping strategies for the majority of the analyses in this study. Strategy 1 was a CPU-based accelerated Sentieon^®^ implementation of the *BWA* ’aln’ algorithm ^116^ and seed lengths of ’-l 16,500’ nucleotides, maximum edit distances set by a fraction ’-n 0.01’ of missing alignments, and ’-o 2’ maximum gap opens ^117^. Strategy 2 was a GPU-accelerated NVIDIA^®^ Clara™ Parabricks ’fq2bam’ implementation of the *BWA* ’mem’ algorithm with settings for short reads (’-M -k 19 -r 2.5’).

Similarly, when mapping ancient reads, we used two strategies for removing duplicate reads and assessing DNA damage. First in the “standard” strategy, we used the *PALEOMIX* ^118^ ’rmdup_collapsed’ function and duplicate removal using *SAMtools* rmdup. We performed base recalibration with *mapDamage* ^119^ to rescale base quality scores in consideration of 12 nucleotides from both 5’ and 3’ ends of sequencing reads (’--rescale-length-5p 12’ and ’--rescale-length-3p 12’). We profiled Ancient DNA damage patterns using DamageProfiler ^120^. The second strategy was “stringent,” where we merged each sequencing run across libraries and marked duplicates again using ’MarkDuplicates’ from the Picard toolkit (<u>broadinstitute.github.io/picard</u>) to remove any remaining duplicate reads at the library level. After additional duplicate removal, we ran mapDamage again to assess overall DNA composition, read length distributions, fragmentation and misincorporation rates and C-to-T and G-to-A transition frequencies at relative positives across reads.

We assessed alignment statistics using *samtools* ’idxstats’ and ’flagstat’, *mosdepth* ^121^, and *QualiMap* ^122^’bamqc’ after removal of marked duplicates: (*i*) *depth of coverage* as the # of unique reads mapping across all reference bases reported as a histogram and the mean # of unique reads mapping, (*ii*) *breadth of coverage* as % bases of reference genome covered by given #s of unique reads reported as a histogram and the % reference at 5× coverage by unique reads, and (*iii*) *physical coverage* or cumulative length of post-filtering and metagenomic classification of collapsed reads expressed as a multiple of the expected 2.8B nucleotides composing the total canid genome size reported from flow cytometry in coyotes, gray wolves, and domestic dogs ^123,124^. As *QualiMap* does not exclude low mapping quality reads, unlike *mosdepth,* alignment statistics may be overestimated.

### Sex inference of ancient specimens

We found support for both DireGB and DireSP individuals as male after mapping libraries to the Greenland gray wolf reference using *BWA* ‘aln’ with stringent duplication removal (see *Read Mapping 1B*). We calculated the Rx ratios ^36,37^ between chromosome X depth and autosomal mean depth of coverage and 95% confidence interval (CI) lower and upper bounds for both individuals, and designated a 95% CI upper bound under 0.6 as male and a 95% CI lower bound over 0.8 as female ^36,37^. The 95% CIs for the ratios of reads aligning to chromosome X versus autosomes placed between 0.539 to 0.554 for DireSP and 0.538 to 0.554 for DireGB.

While the ratios of reads mapping to the reference gray wolf X chromosome (1.0× for SP and 5.7× for GB) and autosomes (2× for SP and 10× for GB) supports these individuals as male, we did not observe similar coverage over the reference gray wolf Y chromosome (0.1× for SP and 0.4× for GB), which may be unsurprising: Y chromosomes are notoriously difficult to assemble given highly repetitive regions and high homology to the X chromosome, and is roughly 1/20th the size of the X chromosome, so mapping sequencing reads from ancient DNA to the Y chromosome may be less stable than mapping to the X chromosome. Therefore, we took a closer look at reads mapping to Y-encoded genes as a secondary confirmation.

We examined depth of coverage by dire wolf reads at the Y-encoded gene *TSPY* because it exhibits relatively high copy number variation among modern gray wolves ^125^ and has high copy numbers in both humans and cattle ^126,127^. Thus, for males we expect to see a relatively higher abundance of reads aligning to *TSPY* than other windows of the assembled Y chromosome sequence. We identified coordinates for the *TSPY* gene annotated on the Greenland gray wolf (*C. lupus*) assembly (GCA_905319855.2) at chrY:594247-596596 (ENSCAFG00865029519 in our gene annotation). We then calculated the depth of coverage by collapsed dire wolf reads aligned using *bedtools* ^128^ and *mosdepth* ^121^ across *TSPY* and 10,000 randomly sampled 2,349 nucleotide windows from assembled chromosome Y and 10,000 windows randomly sampled from any assembled autosome. We observed a high depth of coverage over *TSPY* (0.20× in DireSP and 0.72× in DireGB), relative to depths of coverage from equally-sized 2,349 nucleotide windows randomly sampled 10,000 times over the assembled gray wolf Y chromosome (0.103× ± SD: 0.397× in DireSP and 0.362× ± SD: 1.05× in DireGB), which, while lower than read depth over randomly sampled windows across autosomes (1.87× ± SD: 0.578× in DireSP and 10.6× ± SD: 2.86× in DireGB), supported the presence of male sex chromosomes.

### Multi-canid pangenome construction and read mapping

Ancient DNA mapped to multi-species pangenomes is expected to reduce reference bias from mapping to a single, divergent reference ^89^. Thus we generated a multi-species pangenome of canids using the graph-based *Minigraph-Cactus* (MC) pipeline ^129^, which starts from a reference assembly and iteratively introduces structural variation from aligned genome assemblies and, by excluding centromeric and highly repetitive sequences, can be more efficient for mapping sequencing read data ^130^. We constructed the graph using the Greenland gray wolf (GenBank GCA_905319855.2) as our initial reference genome to be consistent with the coordinate system of our single reference alignments, and incorporated sequence variation from 6 chromosome-level assembled genomes that had the highest contig N50 and most complete set of chromosomes from two *Canis* spp. (domestic dog and dingo) and four other canid species and 3 scaffold-level assemblies from dhole (*Cuon alpinus*), bush dog (*Speothos venaticus*), and red fox (*Vulpes vulpes*) for broad phylogenetic representation of the canid lineage (**Table S2**). We implemented a custom script (’buildPangenome.sh’) launched from the *cactus docker image* (v2.7.1) to construct a pangenome bidirectional sequence variation graph. We built a graph index and mapped unclassified, trimmed reads using *Giraffe* ^131^ included in v*g tools* ^132^. Only reads mapped to the mitochondria (MT) were extracted and utilized for dire wolf MT reconstruction. For downstream analysis, we surjected reads aligned to the multi-canid pangenome graph to the Greenland gray wolf (canLor) reference assembly. We observed that in pangenome alignments by *Giraffe*, short reads were occasionally aggressively clipped, resulting in a small fraction of bases within the overall read aligning. This phenomenon was most pronounced in the dire wolf individuals. We therefore filtered pangenome alignments to remove reads which had more than 20% of their bases soft-clipped, and reads which had a non-clipped length of less than 35 bases.

### Paleogenome reconstruction via iterative mapping and polishing

We reconstructed the dire wolf nuclear genome by iteratively mapping and polishing to a divergent seed reference, the Greenland gray wolf (GenBank GCA_905319855.2), using ancient DNA to progressively assemble a less divergent reference sequence. We performed initial polishing from the DireGB BAM file (see Read Depth) by employing the *Pilon* polisher ^133^ in diploid mode to correct sequences for insertions, deletions, and single nucleotide polymorphisms (SNPs) and perform local reassembly of each chromosome or contig (’--changes --fix all --diploid’).

For subsequent iterations, we allowed each library from DireGB to map to the polished sequences of autosomes, sex chromosomes, and unassembled contigs from the previous iteration, excluding the mitochondrial sequence which we assembled separately. We mapped unclassified and trimmed paired-end reads, merged and unmerged, using the *BWA* ’mem’ and standard deduplication with *PALEOMIX* (see Read Mapping 2A). Unmapped reads were discarded, libraries merged, and sequence polished. We carried out this process for 6 iterations in total and evaluated alignment statistics and quality metrics for each iteration (**Table S4**).

In order to reduce the computational load of assembling the mitochondrial genome and reduce reference bias in mapping, we first mapped the complete libraries of DireGB to the canid Minigraph-Cactus pangenome to identify mitochondrial reads in our data. We surjected all reads that mapped to mitochondrial scaffolds of the canid pangenome to the Greenland gray wolf mitochondrial (MT) sequence and reverse complemented the MT sequence to orient to strand with the D-loop control region. We then filtered reads that were unmerged, unmapped, and not in primary alignment using samtools view -F 261 and removed duplicates with Picard ’MarkDuplicates’ function. The filtered and duplicate-removed mitochondrial BAM was the final input from which we extracted reads using *samtools* ’fastq’ function (-n), resulting in 29,104 reads from the DireGB libraries.

We used *MIA* ^82^, an iterative mapping approach, to assemble the dire wolf mitochondrial genome with the gray wolf genome as the seed reference (NCBI Accession NC_008092.1), giving flags (-c -C -k 14 -U) and -s with an ancient DNA substitution matrix. We filtered the consensus assembly with “relaxed”, requiring 3× coverage and 66% consensus. All sites that did not pass these filters were set to ‘N’. For all downstream analyses, we used the relaxed filtered assembly (**Fig. S5**). We estimated a 34.5× coverage mitochondrial consensus genome for DireSP and a 90.171× coverage mitochondrial genome for DireGB, with 3.35-21.26% N content (**Table S3**).

We performed a final mapping of ancient DNA sequencing libraries to nuclear sequences resulting from the 6th iteration of mapping and polishing plus the iteratively assembled mitochondrial sequence. We aligned using *BWA* ’aln’ with standard *PALEOMIX* deduplication (Read Mapping 1A). We called pseudo-haploid consensus sequences by random read-sampling consensus and by variant-based consensus. In both cases, sequences below 5× unique read depth were rendered as hard-masked Ns, and we required a minimum base quality of BQ=30 and mapping quality of MQ=25. We performed random read-sampling using *ANGSD* ^134^ allele counting (’-doCounts 1 -minQ 30 -minMapQ 25 -dumpCounts 3’) followed by *Consensify* ^135^ to call consensus from reads mapped at 5× to 30× depth and 4/5 allelic support (’-min 5 -max 30 -n_matches 4 -n_random_reads 5’). We called a variant-based consensus to consider read depth and quality scores at variant and invariant sites. We called genotypes using *GATK* ^136^ *HaplotypeCaller* emitting in gVCF mode, then converted to VCF using bcftools convert, filtered variant calls (minimum QUAL=20, BQ=30, MQ=25, DP=5). In the case of overlapping alleles at variant sites, we enforced each genotype to represent homozygous genotype of the allele with highest depth, which should minimize bias from DNA damage-induced cytosine deamination producing false variant calls, using the remove-overlaps plugin for *bcftools* (https://github.com/samtools/bcftools/blob/develop/plugins/remove-overlaps.c). For variable sites, overlapping variants with the highest QUAL values were retained, and for overlapping sites that contain invariant sites, we generated pseudo-QUAL values using the highest QUAL value of a known variant site scaled by the depth of the invariant site prior to filtering. Any sequences not covered in the resulting VCF were hard-masked as Ns in the final consensus.

### Gene annotation and orthology analysis

We annotated genes using the fully automated *FLAG* tool (Troy et al. 2023) implemented in the ’FLAG: Eukaryote Gene Annotation’ workflow on the Form Bio platform (formbio.com). For the reconstructed dire wolf assembly, we included fully assembled transcript data from domestic dog (GCA_000002285.4), gray wolf (GCA_905319855.2), and dingo (GCA_003254725.2) as evidence for *Liftover*, *Augustus*, and *Helixer*. After employing a novel combine and filter algorithm, gene symbols were attributed to each model using *Entap*. We performed a BUSCO analysis using default settings and reported the total number of orthologs found in the mammalian database (n=9226). We assessed mean depth of coverage by all aligned short read libraries over all annotated genes (length-weighted average of mean depth per exon) in the reconstructed dire wolf genome sequence using mosdepth (**Table S5**).

### Consensus calling from iterative reconstruction

We mapped short read genomes of modern canid species sourced from publicly available data sets (**Table S8**), the new sequencing libraries from DireGB and DireSP individuals, and the previously published libraries of dire wolf individuals ^12^ to the reconstructed dire wolf genome reference (aenDir) using *BWA* ’mem’ with standard deduplication with ‘Mark Duplicates’ (Read Mapping 2A).

We called consensus sequences using a random read-sampling approach. We used *ANGSD* (’-doCounts 1 -dumpCounts 3’) to quantify depth of sequence variants with a minimum mapping and sequence quality of 25. We used these counts to generate a consensus genome using Consensify for the modern (’-min 10 -max 500 -n_matches 8 -n_random_reads 10’) and ancient (’-min 5 -max 30 -n_matches 4 -n_random_reads 5’) read sets. Note in both cases, a variable site was incorporated as a change to the reference only if 80% of alternative reads samples were in agreement. Such random sampling of ancient data also decreases the likelihood of incorporating false positive variants from DNA damage signals ^138^.

### *Post hoc* analysis of residual reference bias

We assessed residual reference bias as a disproportionate rate of derived allele sharing with the seed reference Greenland gray wolf (*C. lupus orion* aka “canLor”) genome (GenBank GCA_905319855.2) ^35^ relative to the genome of a distinct gray wolf subspecies (*C. lupus* aka “canLup”). We employed the ABBA/BABA *D*-statistic for introgression ^38,84^ to quantify derived allele sharing. We designated ancestral (A) alleles as those shared with outgroup species gray fox (*Urocyon cinereoargenteus* aka “uroCin”) and/or red fox (*Vulpes vulpes* aka “vulVul”), where for site patterns of derived (D) alleles considered concordant under the expected species tree, ’(((canLor,canLup), aenDir), outgroup/vulVul/uroCin)’ would be ’(((D,D),A),A)’ or ’(((D,D),D),A)’ and discordant as ’(((D,A),D),A)’ and ’(((A,D),D),A)’. We assessed these site patterns using haploid calls from long sequence alignment over syntenic regions to reference genomes of outgroup canid species varying in divergence.

First, we performed a long assembly-to-reference mapping with *minimap2* ^139^ using the preset for up to 10% sequence divergence (’asm10’), and applied the *minimap2* ’paftools call’ function to identify aligned regions (R lines) and sequence variants (V lines) relative to outgroup assemblies from most-to-least diverged from or within the wolf-like clade: the gray fox reference assembly (GenBank GCA_032313775.1) ^112^ and the red fox reference assembly (GenBank GCA_964106925.2) ^111^, the maned wolf (GenBank GCA_028533335.1, *Chrysocyon brachyurus* aka “chrBra”) reference assembly ^107^, and the African wild dog (*Lycaon pictus* aka “lycPic”) reference assembly (Yggdrasil DB12) ^114^. To those reference genomes, we aligned assembled genome sequences of outgroups red fox and gray fox, when not used as reference; the seed-reference Greenland gray wolf (*C. lupus orion* aka canLor) and a non-reference genome assembled for a North American wolf (GenBank GCA_003160815.1, *C. lupus* aka canLup) ^113^; and the pseudo-haploid consensus genome sequences of dire wolf individual DireGB called from random read-sampling ANGSD + Consensify from alignment to either the reconstructed ancient dire wolf (*Aenocyon dirus* aka “aenDir”) genome reference (“iterative”) or its seed-reference sequence of Greenland gray wolf (“divergent”).

We restricted analyses to regions over each outgroup genome aligned by gray wolf and dire wolf genomes, comprising a total of 171,599,977 gray fox aligned sequence bases, 247,756,445 maned wolf aligned bases, and 244,509,933 African wild dog aligned bases, where these taxa should have long-range synteny for sequence alignment and variant calling. We then computed global *D*-statistic as (# ADDA sites - # DADA sites)/(# ADDA sites + # DADA sites) over canid syntenic regions of respective reference assemblies. We computed standard errors via a block jackknife procedure ^38^ to divide the outgroup genomes into 1 megabase equally-spaced contiguous blocks, and adjusted global *D*-statistics by this standard error to obtain Z-scores (**Table S6**).

We mapped DireSP sequence reads using BWA mem as described above to either the iteratively reconstructed dire wolf or gray wolf reference assemblies. Using the *Rsamtools* package (v2.22.0), we sampled 500,000 reads from each alignment file and parsed their cigar strings to calculate the identity-by-state (IBS) for each read. IBS was defined as the (number of aligned bases (M) - number of mismatches (NM))/aligned sequence length (M). Reads were binned by their aligned sequence lengths and average IBS was calculated for each bin.

### Mitochondrial and nuclear phylogenetic analysis

In order to place our newly assembled mitochondrial sequence in phylogenetic context, we downloaded publicly available canid mitochondrial genomes (**Table S9**). We used *rotate* ^140^ to ensure all circularized sequences started at the same position and aligned the sequences with *MUSCLE* v3.8.1 ^141^. We used *RAxML* v8.2.12 ^142^ to build a maximum likelihood phylogeny (-m GTRGamma -f a -# 100 -p 12345 -x 12345) under the GTRGAMMA substitution model with 100 bootstrap replicates. The dire wolf mitochondrial genomes formed a clade outside of other wolf-like canids (**Fig. S5**) in a topology similar to that estimated previously ^12,20,143^.

To localize the placement of the dire wolf in the broader canid phylogeny, we set out to build a nuclear phylogeny using neutral windows of genomic sequence. We defined neutral windows using *bedtools* makewindows (-w 25000 -s 1025000 -i winnum), which defined uniquely named 25 Kb loci at least 1 Mb apart. Windows containing any annotated protein coding genes were removed using bedtools intersect (-v). This resulted in 1,332 possible loci to build trees from.

To assess canid sequence divergence over these windows, we aligned 14 representative canid data sets (**Table S8**), along with DireGB and DireSP, against the iteratively assembled paleogenome reference and obtained consensus sequencing using the *ANGSD* + *Consensify* approach outlined above. Sequences over the neutral windows were extracted and N content was measured using the “letterFrequency” function from the *BioStrings* package in R. Only windows that exhibited <= 50% missingness across all species (n=924) were retained for further analysis. Sequences that passed these thresholds were converted to multiple sequence alignments using *MAFFT* and trees were built from each loci using *IQTree2* with 100 bootstraps each with a GTR + I + G substitution model ^144,145^. Each loci tree was merged and provided as input for *ASTRAL* II ^146,147^ to call a final consensus species tree.

### Divergence time calibration

We used *MCMCtree* v4.10.7 from the *PAML* package ^148^ to perform divergence time estimation. We used an autocorrelated-rates relaxed clock model with approximate likelihood calculation and the HKY85 ^149^ substitution model with gamma-distributed (five categories) rate heterogeneity among sites. We used gamma priors for kappa (shape=2, scale=10) and alpha (1, 10) in the substitution model, and gamma-Dirichlet priors for the overall substitution rate (shape=2, scale=40, concentration=1) and rate drift parameter (2, 20, 1). These priors were selected to provide diffuse yet biologically realistic constraints on the nucleotide transition/transversion ratio, among-site rate variation, overall rate of molecular evolution, and rate drift (allowing for changes in evolutionary rate from parent to offspring branches) in the autocorrelated relaxed clock model, respectively.

We used two nodes as calibration points with priors based on previous studies: the root age calibrated for gray foxes (*Urocyon* spp.) (same root calibration as Perri et al. 2021) versus other canids ^12^ and the South American canid node represented by bush dog and maned wolf ^107^. Lower and upper bounds for the root age were 10.3 Mya (*Metalopex macconnelli*) and 20 Mya (*Leptocyon* spp.), and 1.35-4.34 Mya for the South American canid node. For both nodes, we used log-normal distributions as priors, with 0.025 for location and scale parameters. This approximated a uniform distribution between upper and lower bounds, while softening the bounds with low-probability tails (see prior distributions of node ages in **Fig. S9**). The MCMC analysis was run for 1,000,000 iterations, with the first 10,000 discarded as burn-in, and samples collected every 100 iterations, resulting in 10,000 samples. We created visualizations of the resulting tree with divergence time estimates using the *MCMCtreeR* package ^150^ in R, showing both node age uncertainty and posterior distributions of node ages (**Fig. 3**, **Table S10**).

### Sequence variant calling

We detected germline variants against the reconstructed dire wolf reference (aenDir) from BAMs of 35 modern canids (**Table S8**) and DireGB and DireSP using a joint-calling workflow implemented by *GATK4* ^136^: *HaplotypeCaller* for local re-assembly of haplotypes, *GenomicsDBImport* for gVCF consolidation, and *GenotypeGVCFs* for joint variant calling. We also generated a variant call set comprising transversions only by filtering with *BCFtools* option, ’-i ’((REF="A" & ALT∼"[TC]") | (REF="T" & ALT∼"[AG]") | (REF="C" & ALT∼"[AG]") | (REF="G" & ALT∼"[TC]")) & (strlen(REF)=1 & strlen(ALT)=1)’’, which totaled 41,774,524 SNPs.

We also called variants from the multi-canid pangenome alignments surjected to the Greenland gray wolf reference (canLor) using the GPU-optimized NVIDIA^®^ Clara™ Parabricks implementation of *DeepVariant* ^151^ for 23 modern canids (**Table S8**) and both dire wolves. *DeepVariant* gVCFs were combined using *GLNexus* ^152^ with the ‘DeepVariant_unfiltered’ configuration to avoid biasing allele frequencies of non-gray wolf species. We then set as missing any genotype with GQ<10 using the *bcftools* setGT plugin and filtered a ny site where fewer than 10 individuals were genotyped.

### Sequence distance, diversity, and divergence

We estimated the sequence distance of dire wolves from extant canids using the formula: 2 x (mutations/generation) x (divergence time / generation time), assuming a mutation rate of 4 billion substitutions per generation calibrated for *Canis* species ^41^, a generation time between 5 and 8 years for black-backed jackals and wolf-like canids, respectively ^42^, and a divergence time of 5.8 Mya ^12^.

We assessed sequence similarity in two ways. First, we filtered the final species variant call set (described above) for homozygous transversions by species using *BCFTools* query and compared each using a custom script. Each species in a comparison had to have the same site vary in the same way to be considered a shared allele. We then subtracted the total differences from the non-N sequence length of the reconstructed dire wolf (2,310,286,013 nucleotides) and expressed a proportion of total available sequence. In parallel, we estimated sequence distance using *Mash* ^43^. For each consensus sequence, we used *mash sketch* to hash the genome into 32-mers and assessed common 32-mers in each species in a pairwise manner.

### Gene flow and admixture graph modeling

We converted autosomal variants to *EIGENSTRAT* format using *vcf2eigenstrat* from *SequenceTools* (https://github.com/stschiff/sequenceTools) for both the *GATK4* jointly-called variants against the dire wolf consensus genome and *DeepVariant* variants called against the gray wolf genome from the surjected pangenome alignments. To account for differential coverage across individuals, we then made pseudohaploid genotype calls for each individual for the full set of variants using a read sampling procedure modeled after *Consensify* ^135^ that controls for biases arising from differential sequencing coverage and mapping errors (**Fig. S10, S19**) by randomly selecting 3 reads at each site, or two reads at positions covered by exactly two reads, and requiring at least two bases among the selected reads match. Sites with fewer than two reads were set to missing. This procedure resulted in 33.91M aenDir-mapped and 35.06M pangenome-mapped autosomal, biallelic, mappability-filtered transversions for 17 populations represented by 37 individuals. Ultimately, alignment to the reconstructed dire wolf reference increased reference bias (attraction toward DireGB) in a manner that pangenome alignment did not (**Fig. S12, S19**), particularly for more divergent outgroups. As admixture tests can be particularly sensitive to such biases ^25^, we focused pangenome-derived results for describing canid population history and gene flow, though note that patterns and ratios of admixture in the dire wolf lineage obtained from alignments to the reconstructed dire wolf genome were similar to those inferred from pangenome aligned reads (**Fig. S13**).

We used *D-*statistics to assess allele sharing between dire wolves and extant Canini lineages, using either red foxes or gray foxes as an outgroup *D*-statistics were calculated using *admixtools2* v2.0.6 ^44^. We then used the findgraphs function from *admixtools2* to search for possible well-fitting admixture graph models across varying subsets of canid species. For each set of species, we performed 100 separate iterations of findgraphs across 0-8 admixture events, and extracted the graph with the lowest log-likelihood score as the best-fitting model across each iteration. In cases where less complex graphs provided adequate fits for a given set of species (worst residual of Z< 3), we did not model additional admixture events.

We began with a minimal set of species representing major canid lineages: dire wolves (n=2), Eurasian gray wolves (n=2), coyotes (n=4), dholes (n=2), African wild dogs (n=3), maned wolves (n=2), and red foxes (n=2) as an outgroup. We also explored similar models but with either gray foxes or both red and gray foxes as outgroups. We also explored graphs in which dire wolves were each modeled individually, rather than grouped into a population. Beyond this minimal species set, we also modeled a complex graph which featured African jackals (side-striped jackal and black-backed jackal), along with an additional South American canid lineage (Pampas fox), and a North American gray wolf population (Yellowstone). Finally, we assessed admixture graphs which featured only extant canid lineages and included populations with known history of admixture (golden jackals and red wolves), to assess the sensitivity of this method to detect known instances of admixture among canid lineages.

### Diversifying positive selection analysis

We extracted all protein-coding nucleotide and translated amino acid sequences using *gffread* ^153^ on the pseudo-haploid consensus genome sequences of modern canid species and ancient dire wolf aligned to reconstructed dire wolf paleogenome and called from the random read-sampling *ANGSD* + *Consensify* method, which maintained identical genomic coordinates for gene annotation. We selected the longest coding transcript per distinct gene ID with <20% missing amino acids when translated using *seqkit* ^154^, yielding 13,830 protein-coding genes for positive selection tests. We ran HyPhy analysis programs BUSTED and FitMG94 in local and global modes on aligned CDS and the consensus species tree with dire wolf (DireGB) designated as foreground branch:

(uroCin_SAMN04495241,(vulVul_marble_Fox4,((chrBra_SAMN27257233,speVen_SAMN27257238)1:3.795527799956191,(DireGB,(lupMes_SAMEA5411047,(lycPic_SAMN09924608,(cuoAlp_SAMN10180424,(canRuf_SAMN02921317,(canLup_SAMN02921310,canBai_SAMN02921314)1:0.11623018789473029)1:4.188188635818363)1:0.39859685674669815)1:0.323225231778891)1:0.1826891919310381)1:1.4874594855476162)1:6.424328629404271):0.0);

We tested protein-coding genes using the BUSTED model for episodic positive selection to identify where at least one site experienced positive selection in the dire wolf branch. Additionally, we extracted from the FitMG94 output the branch-specific dN/dS rate estimates under a free-ratio model (’--type local’) and respective non-synonymous and synonymous branch lengths for dire wolf and 10 other canids and the likelihood-ratio test for overall dN/dS != 1 across the entire species tree using a one-ratio model (’--type global’). We considered those genes with tree-wide, one-ratio dN/dS estimate > 2 as indicating pervasive positive selection over the canid lineage.

We flagged genes as under episodic positive aka diversifying selection in the dire wolf branch that exhibited *p* < 0.05 in the BUSTED test and which flagged at least a single codon of evidence ratio > 2, and excluding genes for which the dire wolf consensus codon did not disagree with dire wolf reconstructed reference codon and neither the focal dire wolf codon nor root canid codon contained missing bases (Ns). The process of consensus calling at sites of low depth (<5×) or heterozygous alleles introduces missing bases that could result in misassignment of codons for the focal branch (e.g., dire wolf) and/or root branch (ancestral canid), which would in turn lead to false positives as HyPhy BUSTED needs only a single site to reject fitting the constrained model (no positive selection). Furthermore, BUSTED is insensitive to synonymous substitution rate variation (a constant dS=1 is held in all cases).

We also flagged additional genes as having strong evidence for positive aka diversifying selection in the dire wolf branch, if these had free-ratio maximum likelihood estimates of dN/dS > 2 from the standard MG94 codon evolution model ^155^ implemented in the HyPhy FitMG94 program. We filtered out such genes depleted of synonymous substitutions where dS > 0.001, to avoid extreme estimates, and no excess saturation of substitutions where dS or dN was greater than 4.0, and which exhibited overall dN/dS estimated under the one-ratio model meeting statistical significance (α=0.05) in the log-ratio test given degrees of freedom=(estimated parameters in the standard MG94 model - estimated parameters in the nucleotide GTR model) > 0 under the chi-squared distribution. We excluded those that exhibited extremely saturated rates of substitutions (dS or dN > 4.0) or depleted rates of synonymous substitutions (dS < 0.001), because dN/dS ratios could be inappropriately estimated or result from misalignment of reads ^156^.

We report all protein-coding genes tested and respective statistics from HyPhy BUSTED and FitMG94 programs in **Table S11**. As BUSTED evidence ratios only indicate whether a site better fits under the alternative, unconstrained model that permits positive selection, rather than providing direct evidence site-specific positive selection, we also ran HyPhy FEL (fixed effects indicating pervasive selection) and MEME (mixed effects aka partitioned selection) maximum likelihood models over all 80 genes under diversifying positive selection in dire wolves.

We performed gene set enrichment analysis using *g:Profiler* (’gprofiler2`’)` ^157^ and the domestic dog ortholog reference set (organism: “clfamiliaris”), testing the combined suite of 80 genes under episodic diversifying selection against the background set of 5,497 genes tested that pass all aforementioned filters. Corrections for multiple testing were performed using the Benjamini-Hochberg procedure.

We contextualized dire wolf specific substitutions using *InterProScan* to predict protein functional domains ^53^ and ESM1b model to predict substitution impact via *in silico* mutagenesis ^54^ over each dire wolf protein sequence. We replaced all invalid amino acid residues (’X’ introduced from N content in coding sequences) with a neutral amino acid, glycine (’G’), to enable modeling. For protein sequences longer than 1,022 amino acids that hit the sequence limit of the ESM1b model, we performed tiled predictions as in Brandes et al. 2023 ^54^ to obtain weighted average effect scores. We examined residues harboring dire wolf specific amino acids and compared to the canid-majority amino acid at each residue. As the “wild type” sequence modeled is sourced from the reconstructed dire wolf reference (aenDir), we considered the positive or negative pseudo log-likelihood scores between the consensus amino acid assigned to the dire wolf branch in BUSTED and the canid majority amino acid, depending which amino acid matched the root in BUSTED analysis. We report domain and effect predictions for flagged residues modeled in **Table S12**.

## Supporting information

HyPhy results

## DATA AVAILABILITY

The sequencing data underlying this study are available in the NCBI Sequence Read Archive, and can be accessed with BioProject accession PRJNA1222369. For the HyPhy analyses, we included a tarball of the multiple sequence alignment FASTAs of the coding sequences for all 80 genes under diversifying selection in the dire wolf branch as well as the HyPhy results JSON files from BUSTED, FitMG94, FEL, and MEME programs. These data and all supplemental tables are available in a Zenodo repository associated with this study under the BioRxiv DOI.

## ACKNOWLEDGEMENTS

We thank the Center for Genome Innovation and the Computational Biology Core in the Institute for Systems Genomics at the University of Connecticut for sequencing and data management support. We thank the Bureau of Reclamation for sampling permissions of the Idaho individual that produced DireGB (IMNH 48001/52) and staff at the Idaho Museum of Natural History for supporting resampling of specimens under their care. We thank the staff including Glenn Storrs and Cameron Schwalbach of the Cincinnati Museum Center for additional sampling of the DireSP specimen (VP1737). We also thank colleagues who lent their time and expertise to provide valuable comments on this work.

## AUTHOR CONTRIBUTIONS

Conceptualization: G.G., K.M.P, J.O., M.CJ., B.L.C, B.S.

Data curation: G.G., K.M.P, J.O., B.L.C.

Formal analysis: G.G., K.M.P, J.O., C.H., M.CJ., W.T.

Funding acquisition: B.L.

Investigation: S.B., C.C., J.M., B.R.P, P.G.S.G., S.J.H., S.S., W.S., R.J.O, W.T.

Methodology: G.G., K.M.P, J.O., M.CJ., B.L.C, B.S.

Project administration: G.G., K.M.P

Resources: N.A, S.B., B.R.P., J.M., S.J.H., R.J.O, S.S., W.S., C.C., L.K., T.M., B.S.

Software: N.A., G.G., K.M.P, J.O., M.CJ.,

Supervision: B.L.C, B.S.

Validation: G.G., K.M.P., J.O.

Visualization: G.G., K.M.P, J.O., M.CJ, C.H.

Writing – original draft: K.M.P., G.G.

Writing – review & editing: G.G., K.M.P., J.O. M.CJ., C.C., N.A, W.T, O.F, S.B., T.M., L.K., J.K., A.D., M.C., J.B.P, S.O., R.J.O, S.J.H., P.G.S.G., S.S., W.S., J.M., B.R.P, G.L., L.F, A.P., A.T.L, M.S.S., M.T.P.G, S.G., B.vH., E.K.K., C.E.M., L.D, B.L.C., G.R.R.M., G.C, B.S.

## SUPPLEMENTAL INFORMATION

Document S1: Figures S1-19

Table S1: Sequencing library statistics for all ancient samples.

Table S2: Reference assembly accessions used in study

Table S3: Alignment statistics for all ancient samples

Table S4: Iterative mapping and polishing statistics for paleogenome reconstruction

Table S5: Gene annotation information for paleogenome reconstruction

Table S6: Allele sharing statistics for residual reference bias assessment Table S7: Genetic distance metrics

Table S8: Short read dataset accessions used in study

Table S9: Mitochondrial genome accessions used in study

Table S10: Age estimates for nodes in phylogenetic tree

Table S11: Summary of admixture methods and models

Table S12: Selection test results by gene

Table S13: Genes with dire wolf specific substitutions

