## Supplemental Figures S1-19 for "On the ancestry and evolution of the extinct dire wolf"

### Supplementary Figures

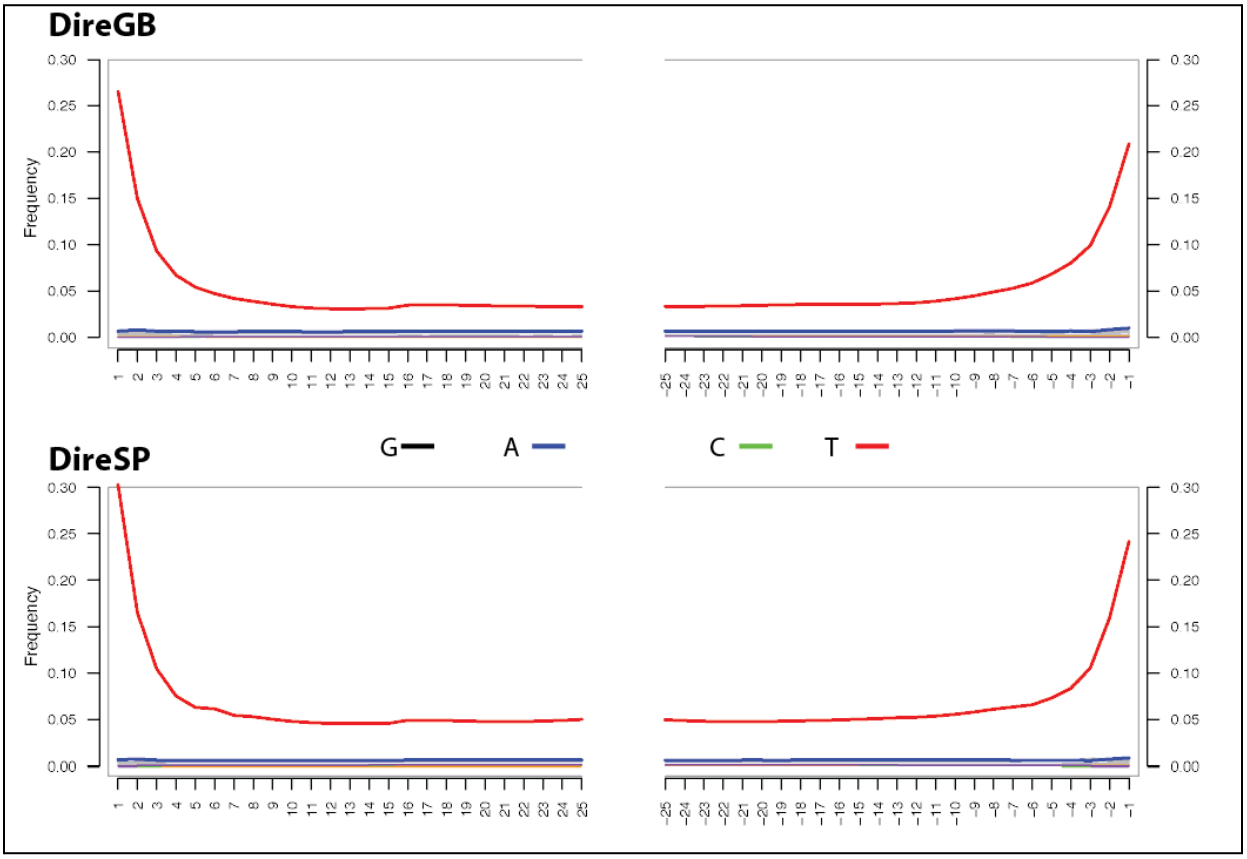


**Figure S1.** DNA damage profile of libraries from two *Aenocyon dirus* specimens, DireGB and DireSP. Both libraries exhibit increased cytosine deamination at both ends of molecules, typical of single-stranded libraries.

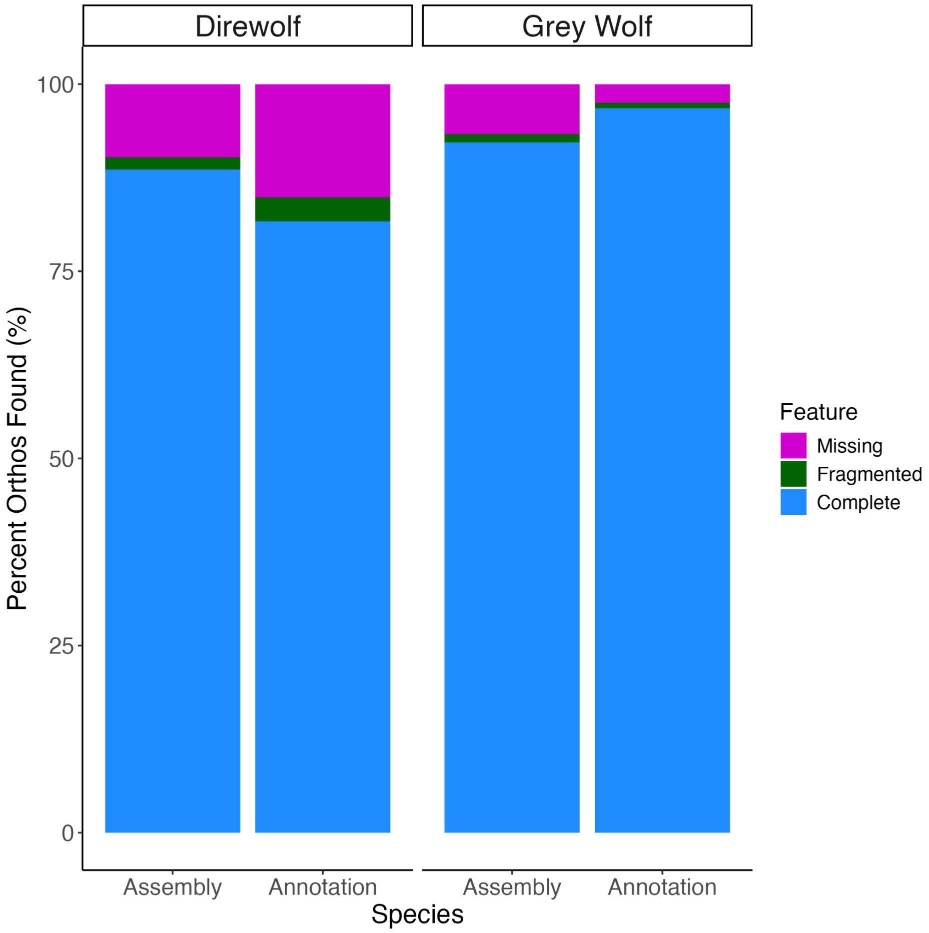


**Figure S2.** Completeness measures of ancient annotation. BUSCO statistics of mammalian orthologs for the dire wolf reconstructed genome assembly and annotation produced using FLAG. Grey wolf measurements reported from Ensembl. The majority of single copy protein orthologs identified in the dire wolf assembly are captured by the FLAG annotation pipeline.

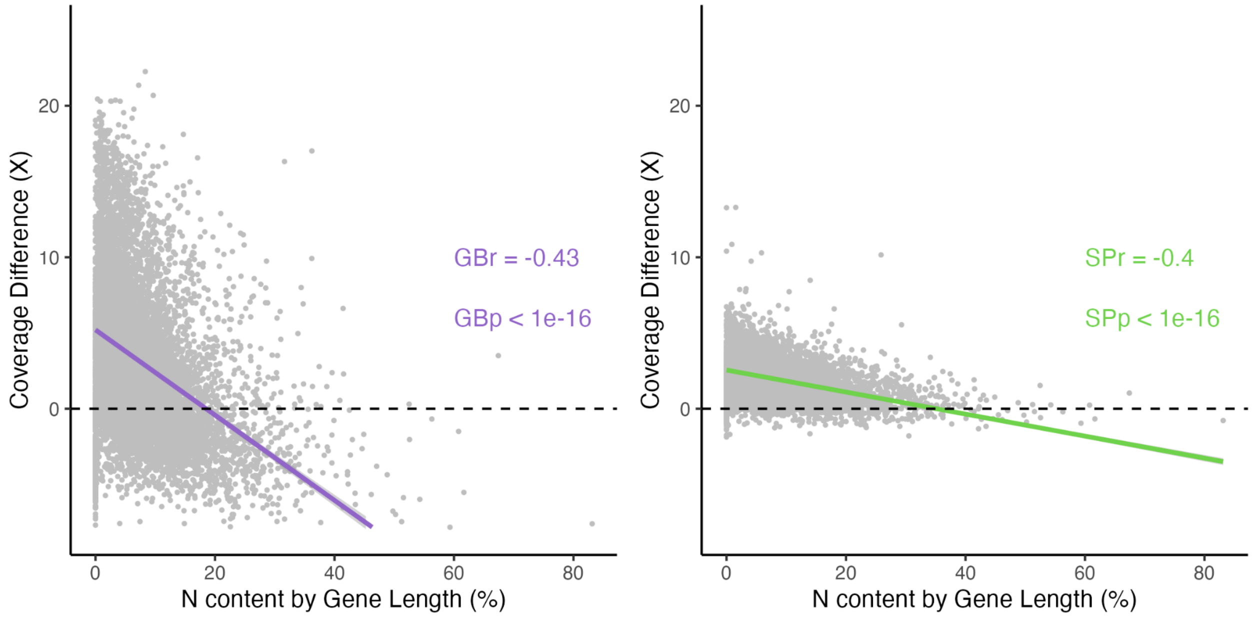


**Figure S3.** Decreased coverage over reconstructed dire wolf exome is correlated with missing data. The total coverage difference when mapping either DireGB (purple) or DireSP (green) to either the reconstructed dire wolf reference or the gray wolf reference. Positive values indicate improved coverage with dire wolf reference. N content is measured as the total number of N bases across the full CDS of a given gene. Only genes confidently annotated in both assemblies (n = ?) were used for comparison. Correlation test conducted in R.

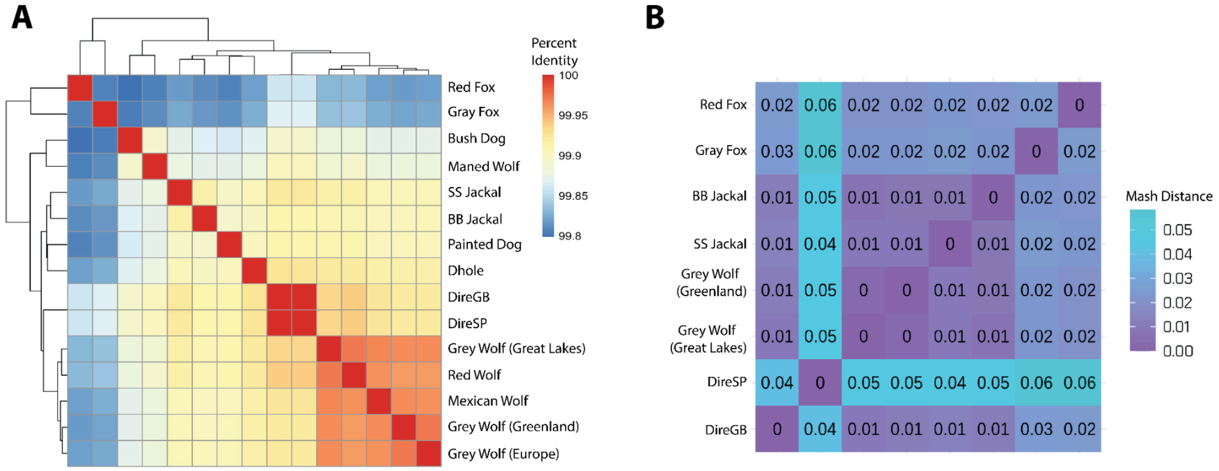


**Figure S4.** Sequence similarity across canid genomes. **A**) Heatmap showing pairwise percent similarity across a variety of canid species. Percent similarity was determined by comparing the number of shared homozygous transversions over the total non-N proportion of dire wolf reference sequence (~2.3Gb). Warmer colors indicate greater sequence similarity (i.e. fewer shared variants). **B**) MASH distance comparing common sets of 32-mers in foxes, jackals, wolves, and dire wolves (n = 2 each). Warmer colors indicate more shared kmers and greater sequence similarity.

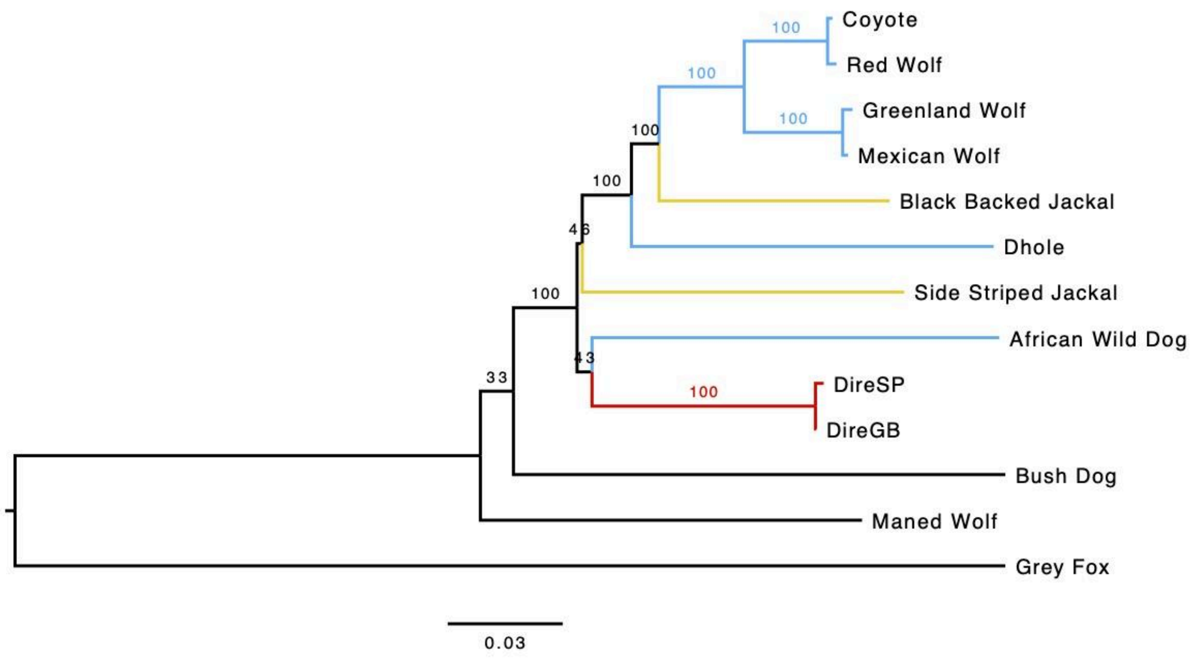


**Figure S5.** Phylogeny of dire wolf mitochondrial DNA. Consensus tree built from full length mitochondrial sequences from wolf-like canids (blue), jackals (yellow), dire wolves (red) and other canids (black). Bootstrap support values (n = 100 replicates) from RAxML are displayed for each node.

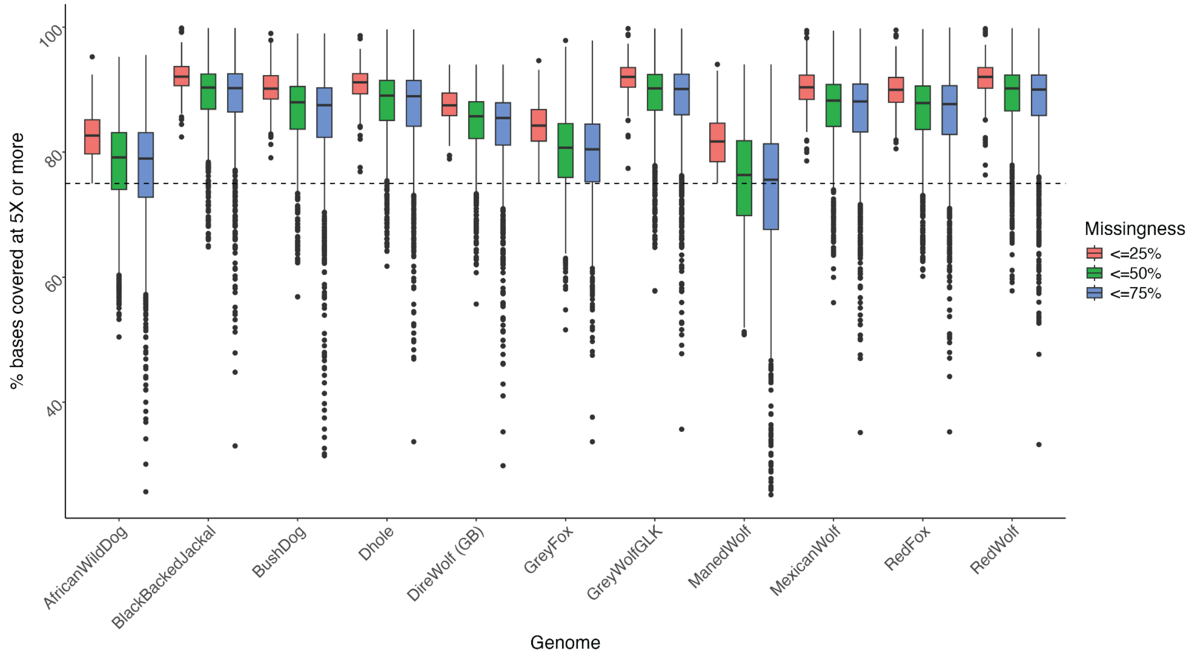

**Figure S6.** Missing data in phylogenetic sequence windows across species. Boxplots display distribution of 25kb neutral windows at either 25% (red), 50% (green), or 75% (blue) maximum missingness thresholds. Dotted line indicates windows with 75% of bases covered at 5X or more (i.e. 25% missingness). While additional windows are added as the missingness threshold increases, missingness does not surpass 25% on average across all thresholds and species.

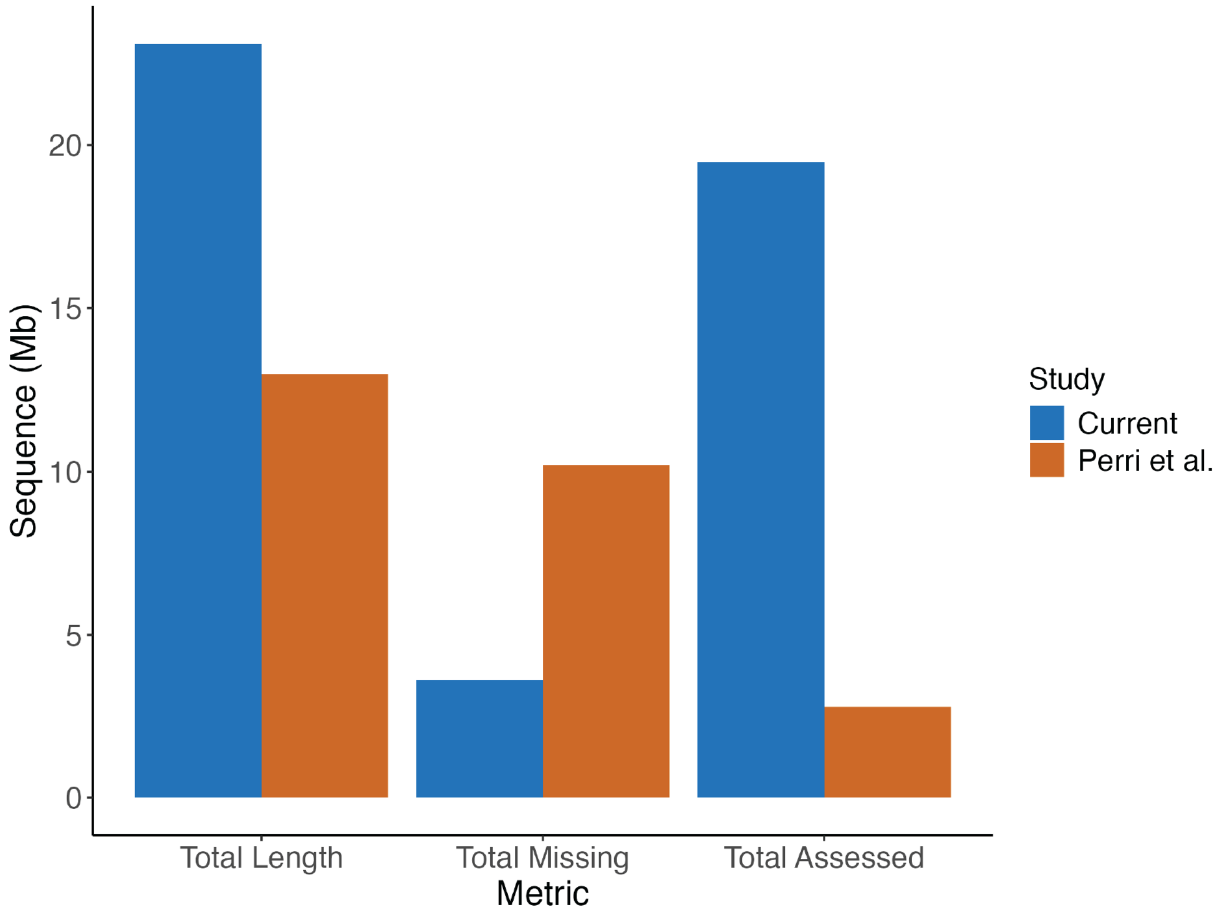


**Figure S7.** Alignment statistics for DireGB data utilized in nuclear phylogeny. Total aligned bases of DireGB across all windows of the alignment. For the present study, total alignment length was calculated by summing all 25kb windows used (n=924), missingness was determined by summing all N bases over these windows in DireGB, and total assessed was the difference. All values were compared to those reported in Perri et al. (2021) for the analysis using DireGB only.

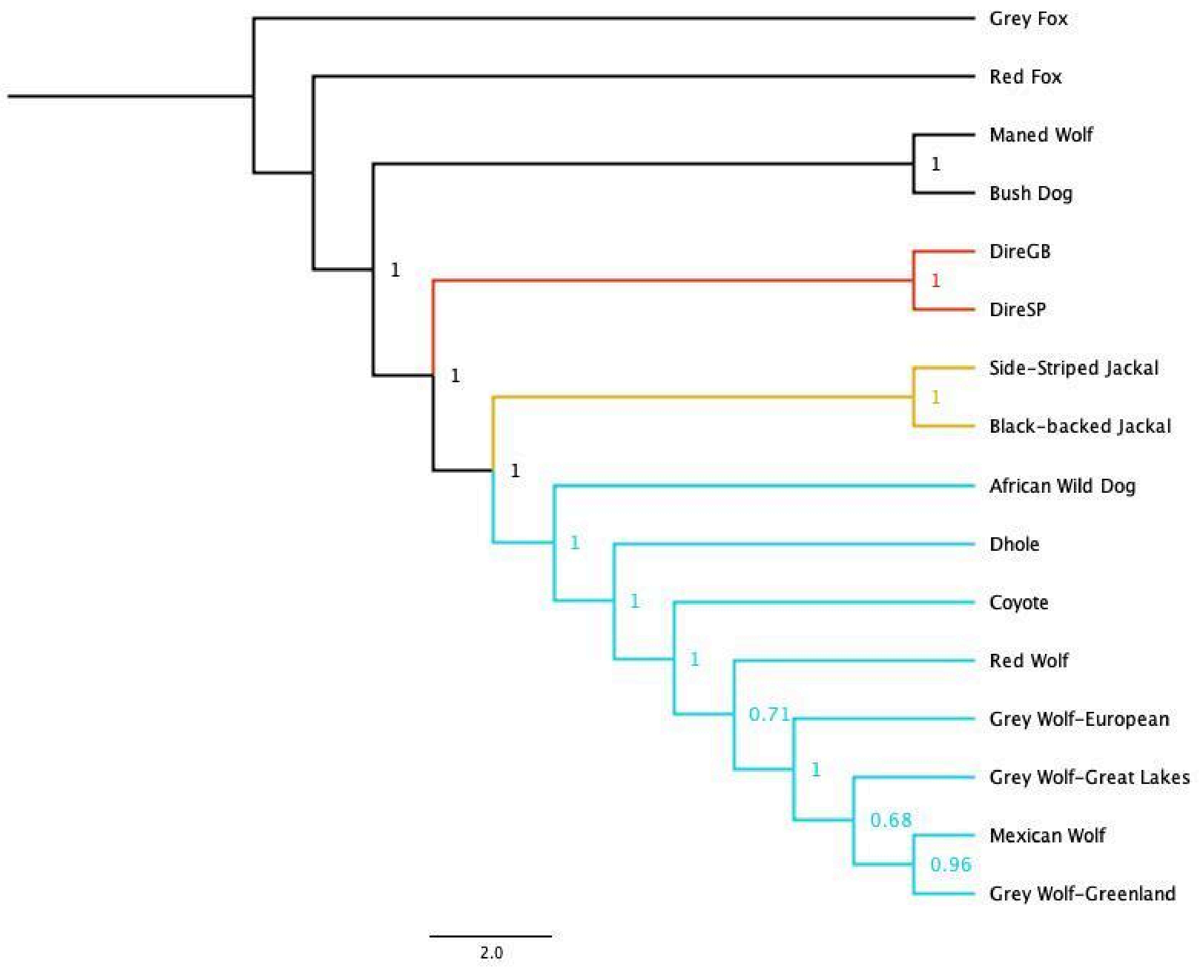


**Figure S8.** Nuclear phylogeny with both specimens confirms monophyletic placement of dire wolf branch. Nuclear phylogenetic relationship of dire wolves (DireGB and DireSP) from 25kb loci trees (n = 861) with no more than 75% missingness across all windows in all samples. Branch lengths presented in coalescence units and node labels are posterior probabilities from consensus tree calling from ASTRAL from each bootstrapped loci tree (n=100 replicates each). Dire wolves (red) form a monophyletic clade branching prior to the split of jackals (yellow) and wolf-like canids (blue).

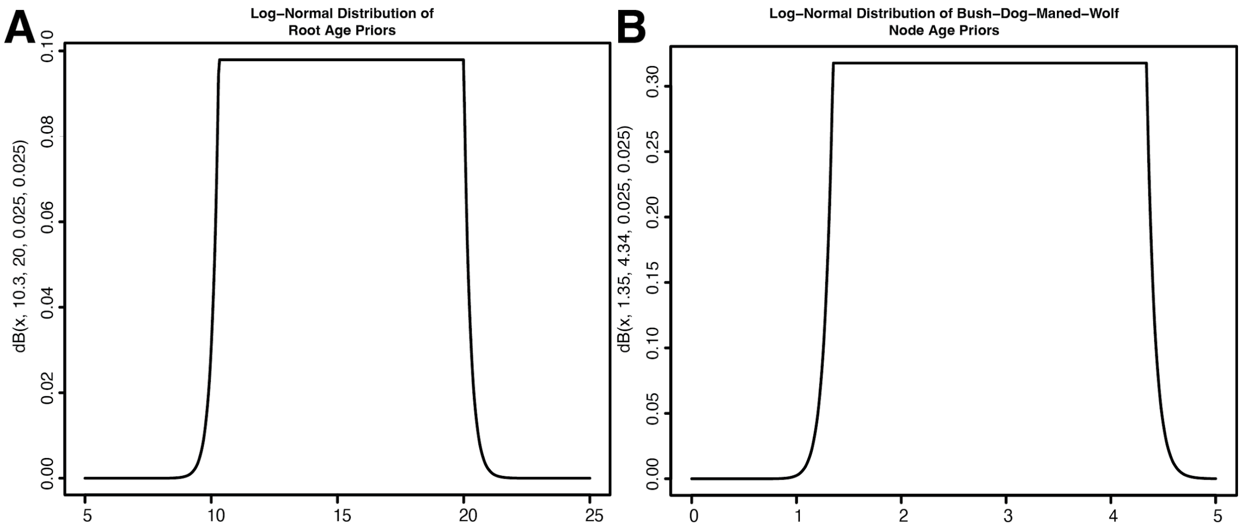


**Figure S9**: Log-normal distributions of age priors for two key nodes in the phylogenetic tree of canids. **A)** Root age prior, representing the split between gray foxes *(Urocyon* spp.) and other canids, with bounds of 10.3-20.0 million years ago (mya) based on Perri et al. (2021). **B)** South American canid node prior, representing the divergence of bush dog and maned wolf, with bounds of 1.35-4.34 mya derived from Chavez et al. (2022). These priors were designed to approximate uniform distributions within specified bounds while allowing for low-probability tails beyond these limits.

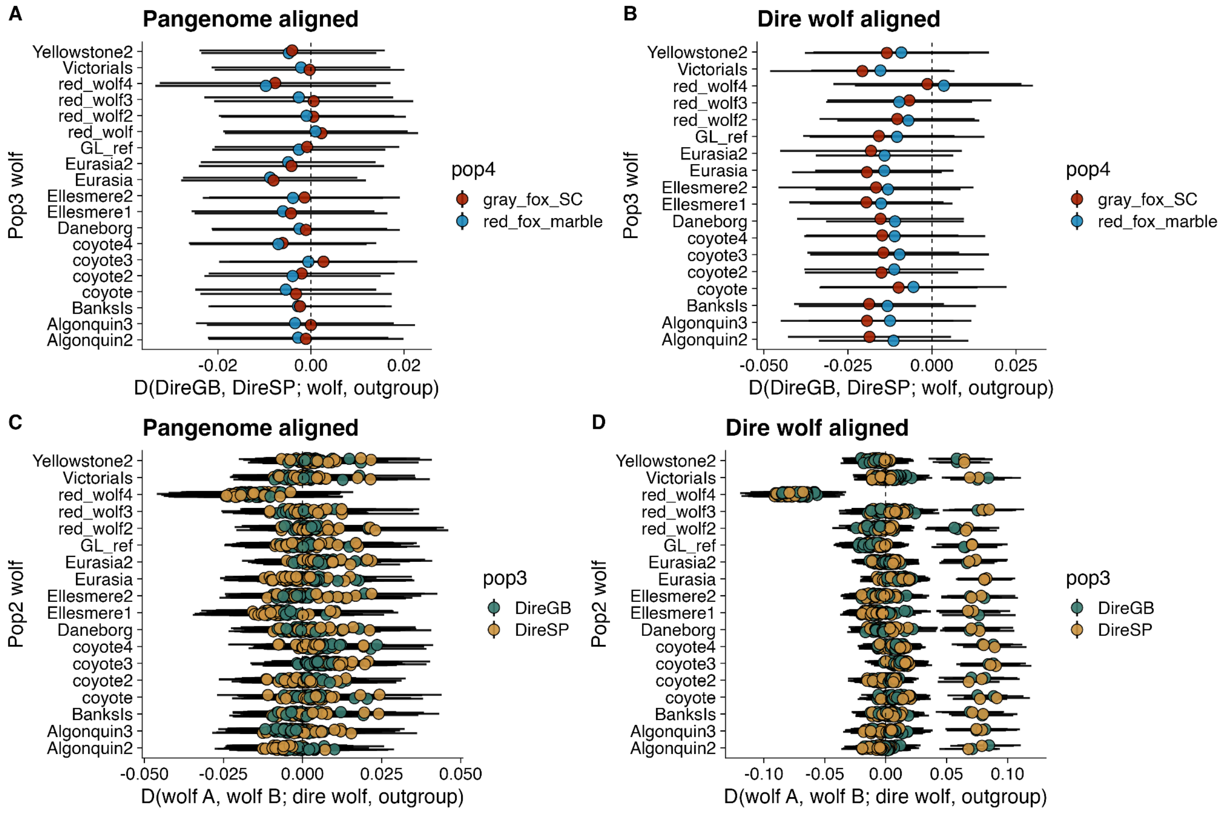


**Figure S10.** *D*-statistics testing whether the two dire wolf specimens, DireGB and DireSP, form a clade relative to different individual wolves, coyotes and red wolves **(A,B)**, and whether all wolves, coyotes, and red wolves for a clade relative to each dire wolf specimen **(C, D)**. **A)** and **C)** show *D*-statsitics calculated using Consensify pseudohaploid genotypes from alignments to a canid pangenome, while **B)** and **D)** show *D*-statsitics calculated using Consensify pseudohaploid genotypes from alignments to an iteratively polished dire wolf reference.

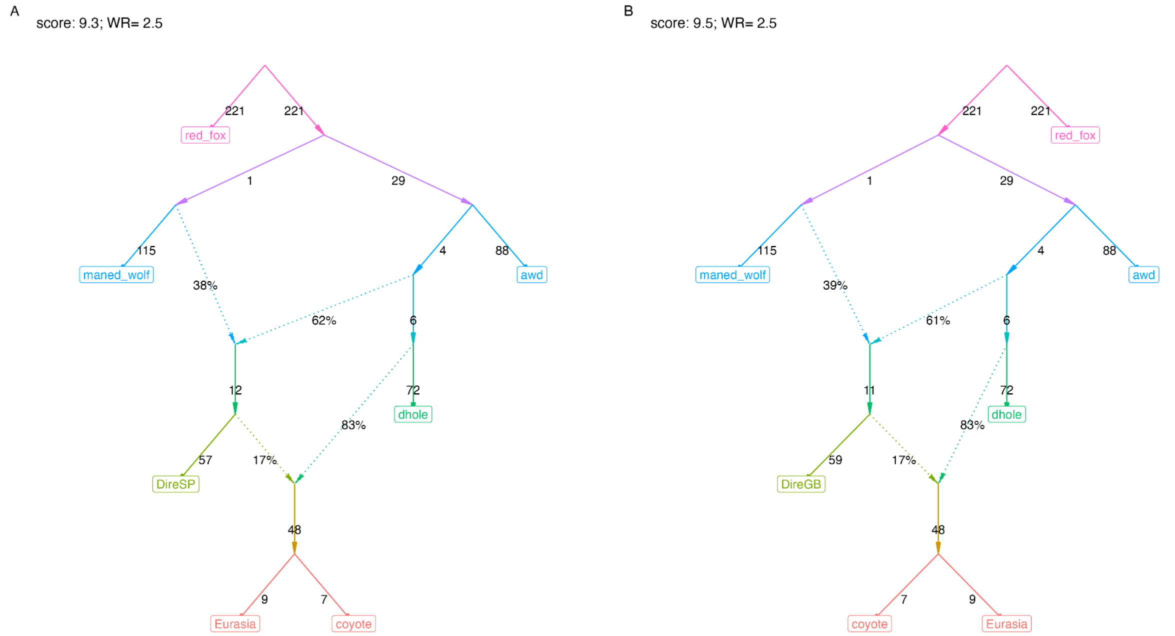


**Figure S11.** Best-fitting admixture graphs for DireSP (**A**) and DireGB (**B**), with the equivalent model grouping both dire wolf individuals presented in Fig. 4C.

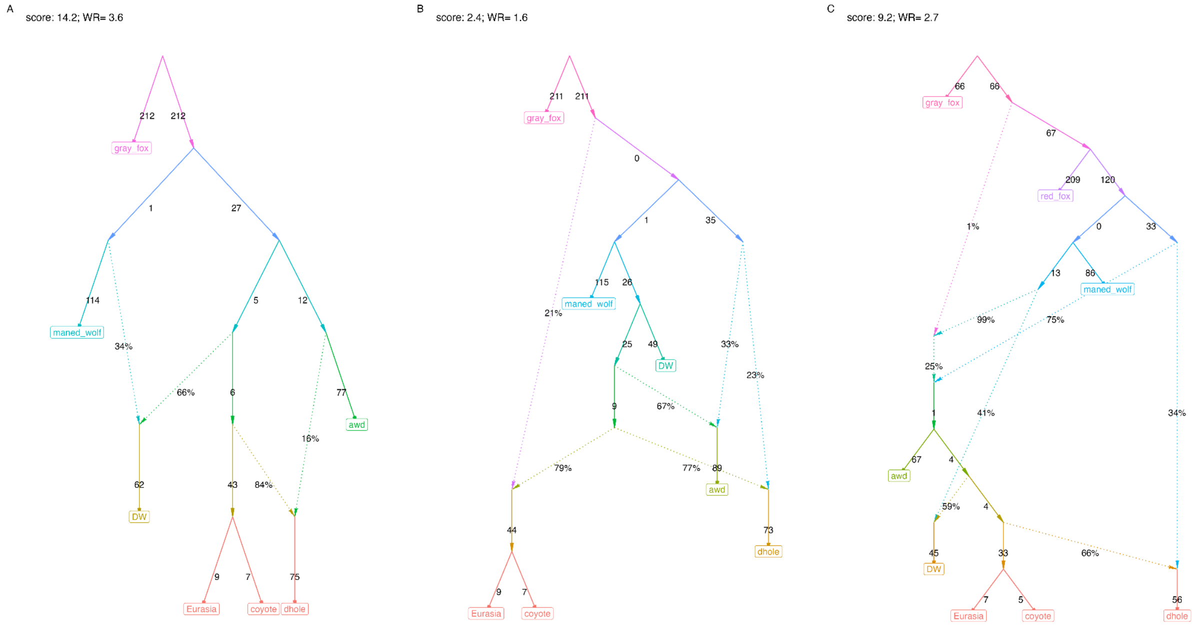

**Figure S12** Best-fitting admixture graphs with gray fox as an outgroup, fit with either **A)** two or **B)** three admixture events, and **C)** with the inclusion of red fox, a closer outgroup. Graphs were fit using Consensify pseudohaploid genotypes called from alignments to the canid pangenome.

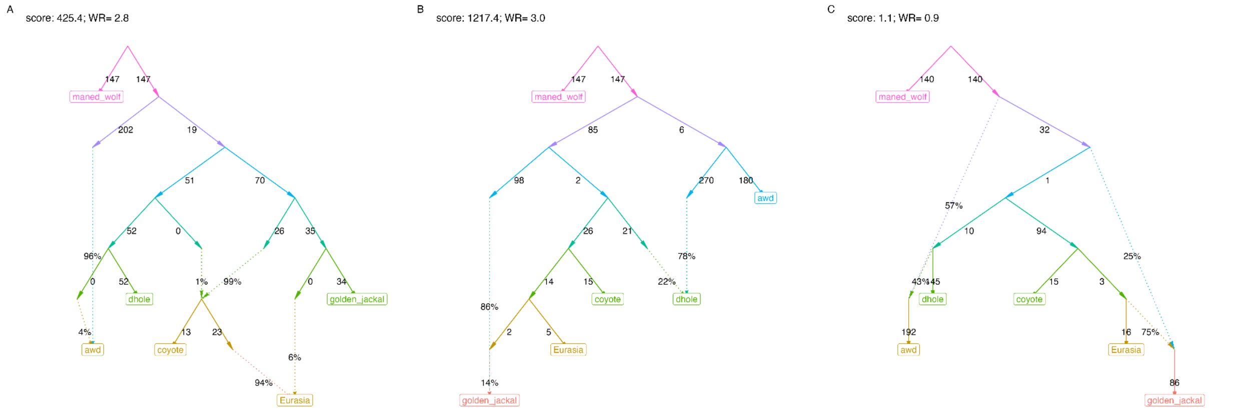


**Figure S13.** Admixture graphs investigating an excess of basal ancestry in wolves and coyotes, relative to dholes and golden jackals. **A**) Graph depicting a "ghost" lineage related to dholes which contributed ancestry to wolves and coyotes. **B**) Alternative graph in which golden jackals and dholes are both admixed. **C**) An alternative graph discovered using *findgraphs* in which African wild dogs and golden jackals are admixed. While all three graphs provide reasonable fits to the data (worst residual < 3), the graph presented in **C**) is the best-fitting and does not necessitate gene flow from a "ghost" lineage into the ancestor of wolves and coyotes.

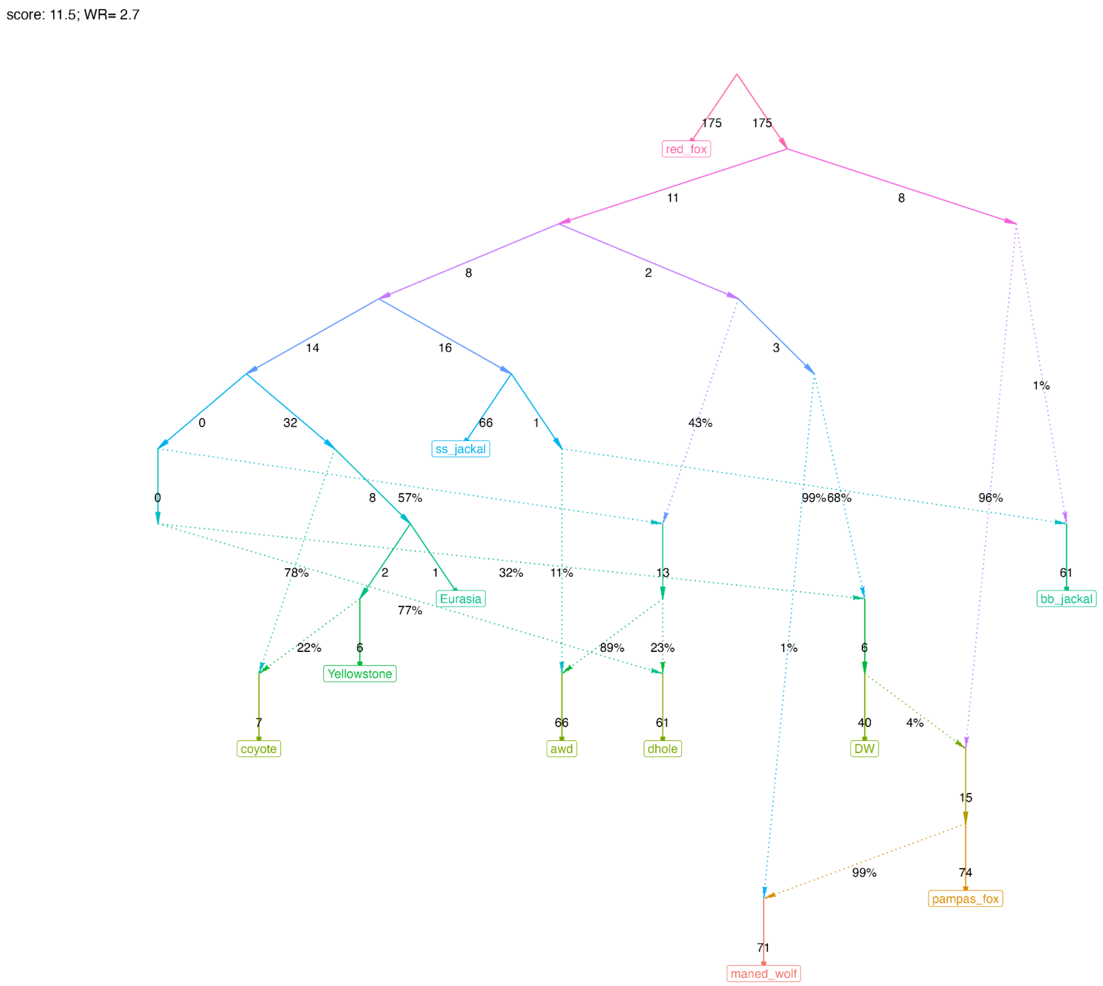


**Figure S14.** Best-fitting admixture graph model with 8 admixture events modeling population relationships across Canini.

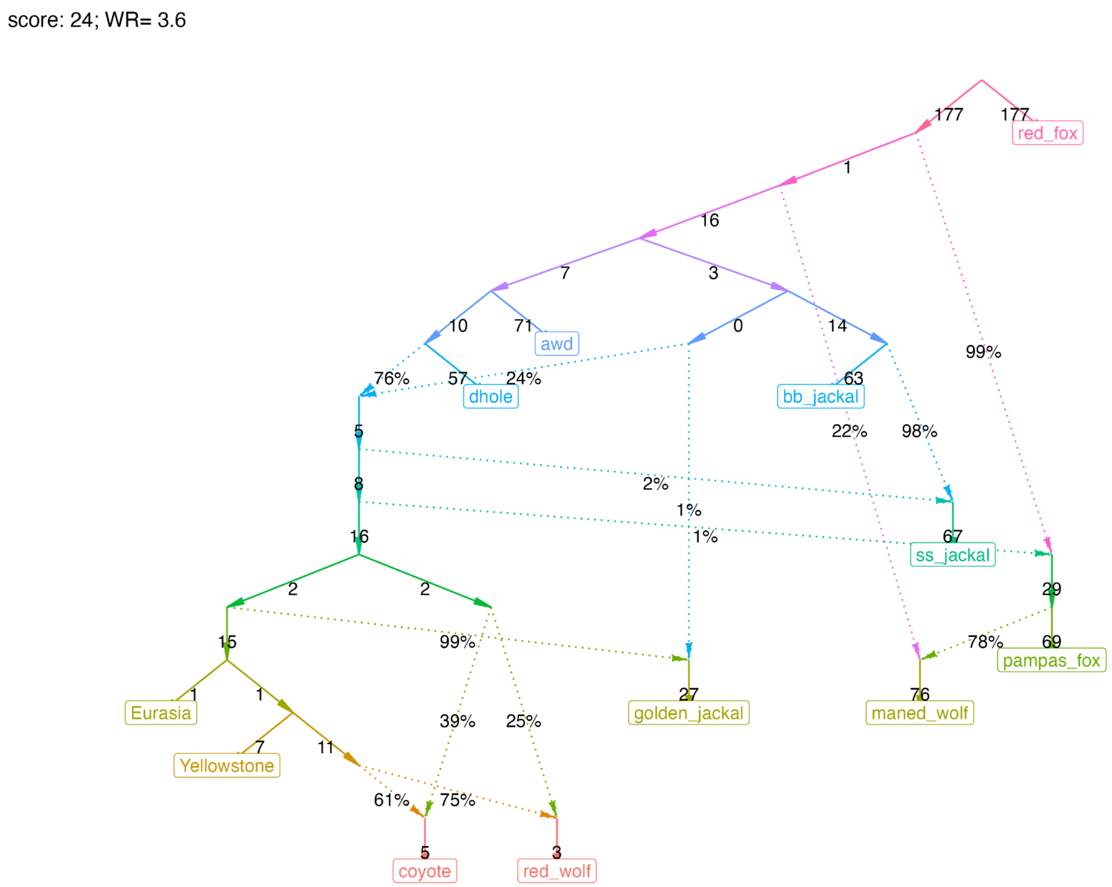


**Figure S15.** Best-fitting admixture graph with 7 admixture events which recovers gene flow events in species with known histories of admixture (including North American gray wolf ancestry in coyotes and red wolves, and gene flow between gray wolves and golden jackals).

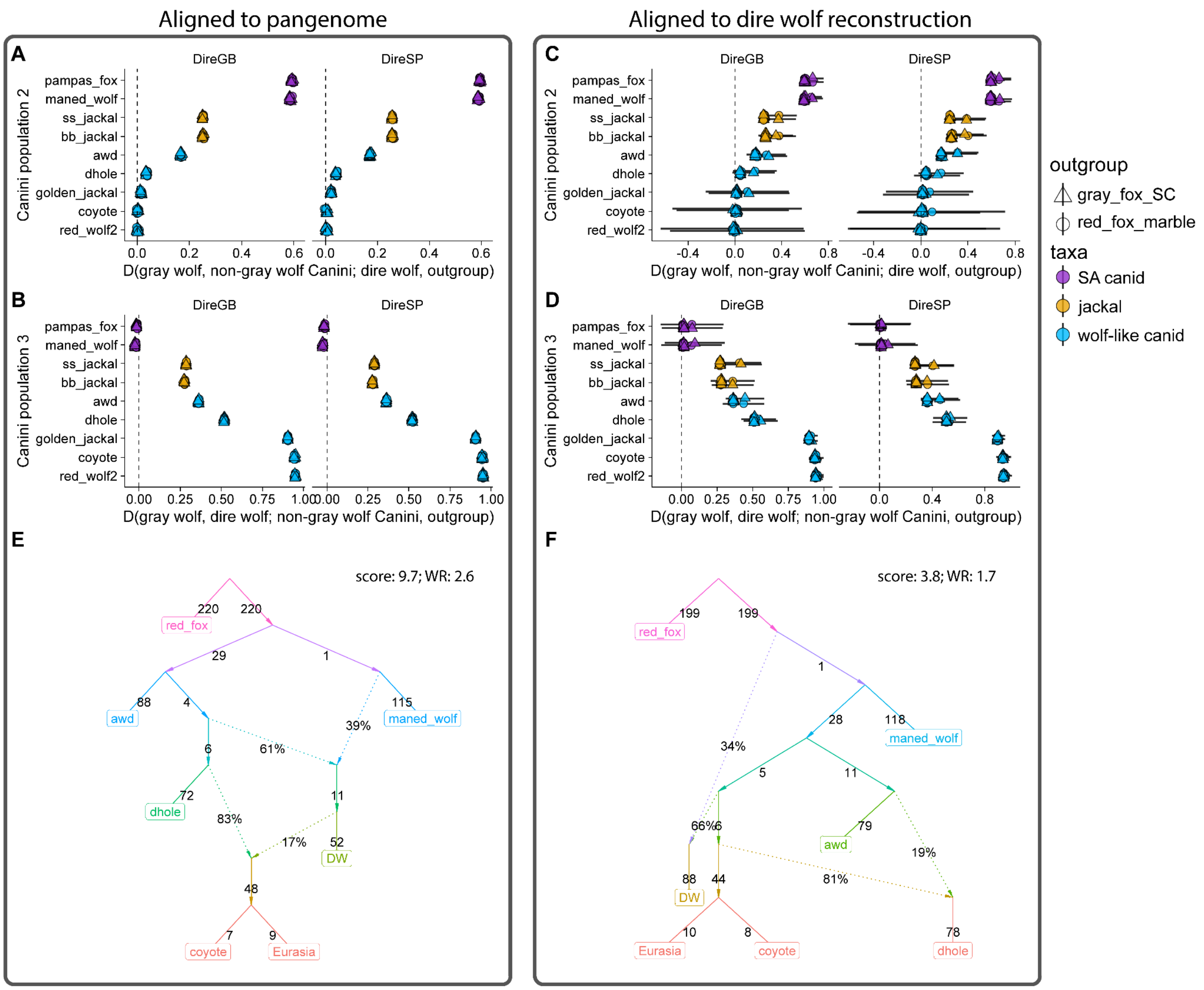


**Figure S16.** Comparison of admixture analyses using either a canid pangenome (**A**,**B**,**E**) or the dire wolf iterative genome (**C**, **D**, **F**) as a reference. **A**) and **C**) show allele sharing between dire wolves and gray wolves, relative to other species within Canini. **B**) and **D**) show allele sharing between gray wolves and other Canini species, relative to dire wolves. **E**) and **F**) depict the best-fitting (lowest log-likelihood score) admixture graphs with two admixture events modeling the relationship between dire wolves and other canids. Both graphs model dire wolves with a mixture of basal and wolf-like canid ancestries, though the phylogenetic placement of the basal ancestry differs. The pangenome graph depicts the ancestors of coyotes and gray wolves as being admixed, while the dire wolf genome reference instead models dholes as admixed.

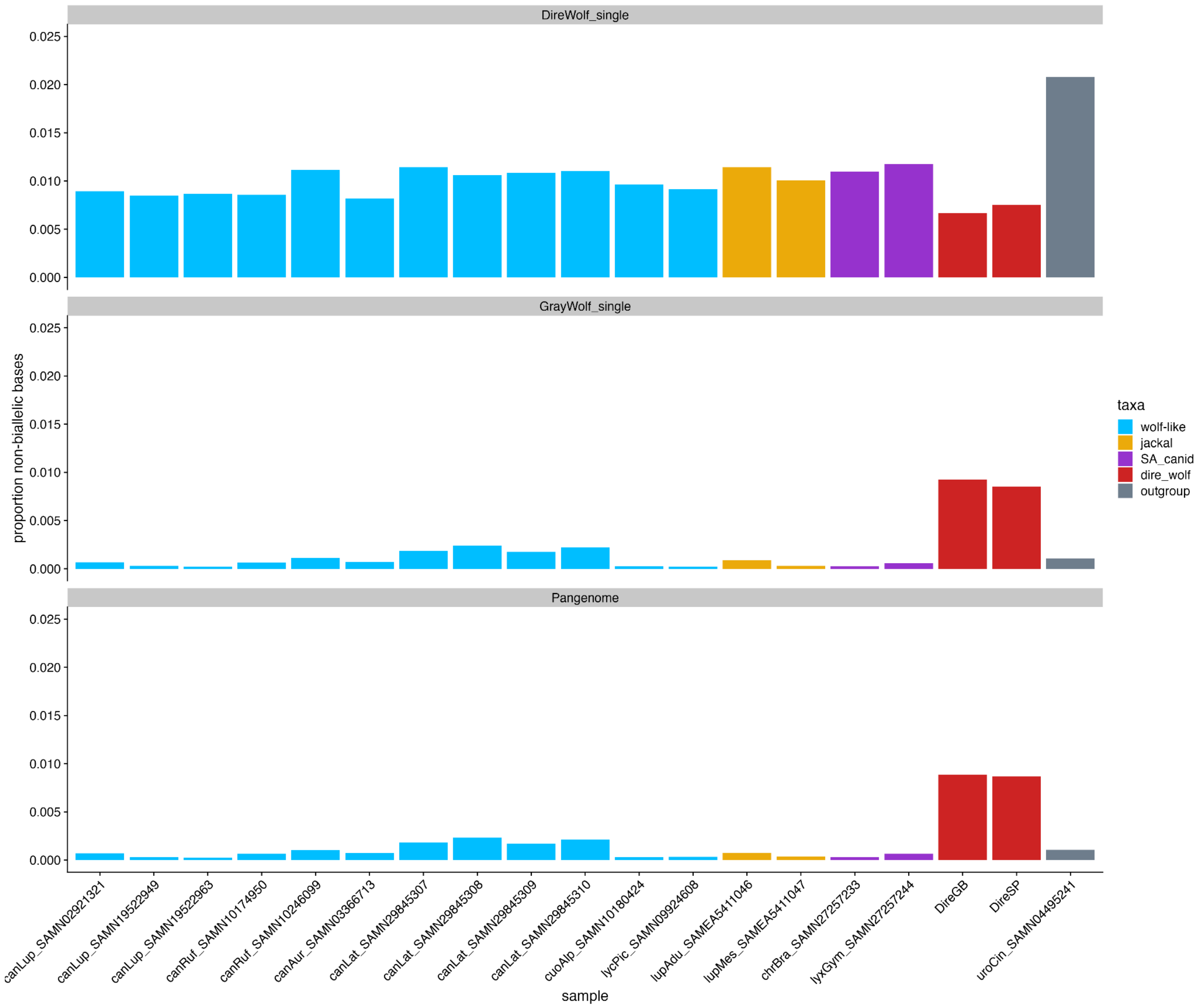


**Figure S17.** Rate of non-bialleleic bases occurring in aligned reads at sites determined to be biallelic across canids. This is a proxy for relative error rate and reflects the rate of unexpected bases occurring due to either sequencing or mapping error, or ancient DNA damage. The rate of unexpected bases occurring is shown for reads mapped to the iteratively polished dire wolf genome (top), the gray wolf reference genome (mCanLor; middle), and a canid pangenome (bottom).

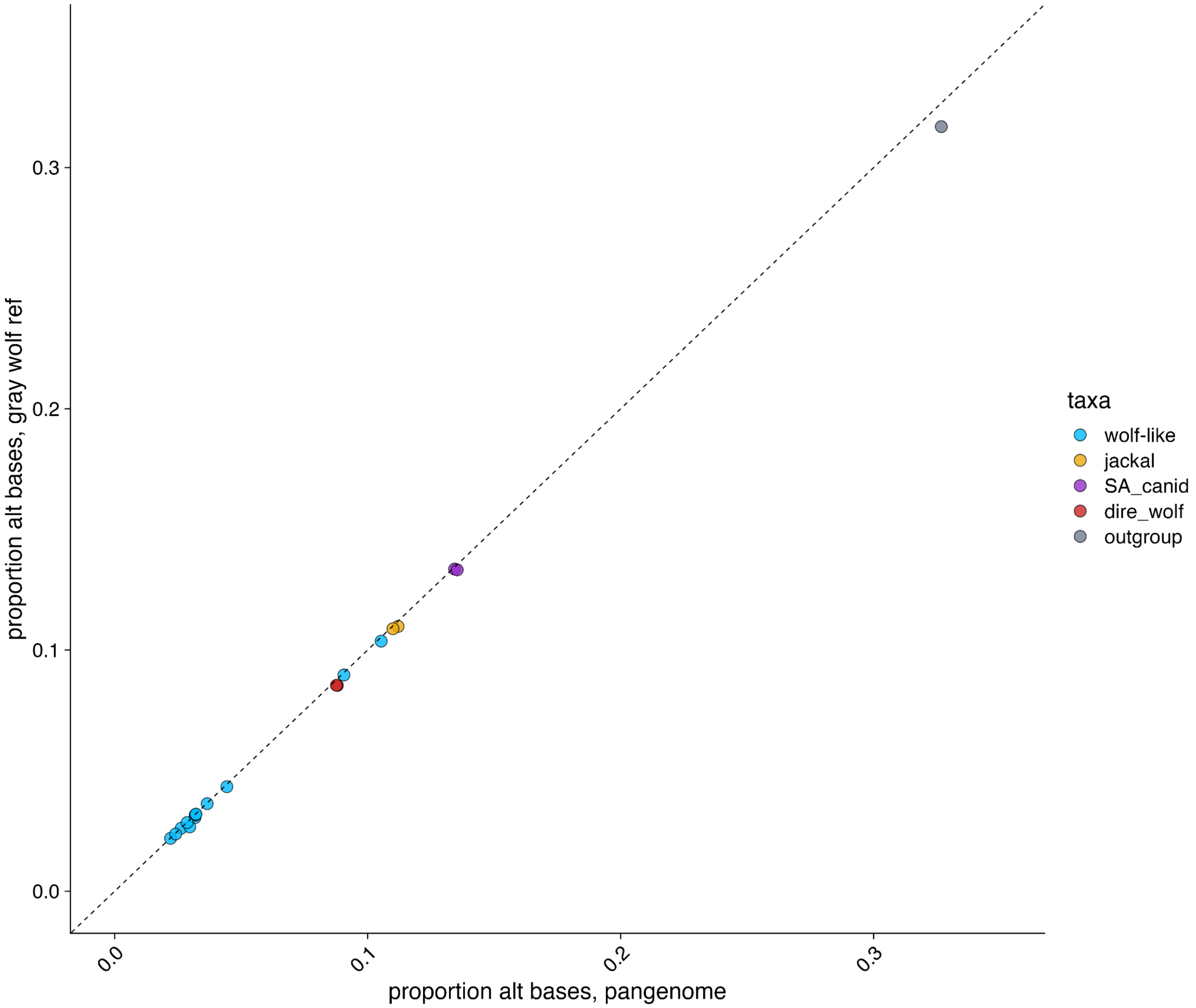


**Figure S18.** Proportion of bases carrying the alternative allele at biallelic sites for all canid individuals aligned either to a canid pangenome (x-axis) or to the gray wolf reference genome (y-axis). Pangenome alignments have a greater proportion of alternative alleles, especially for more divergent taxa.

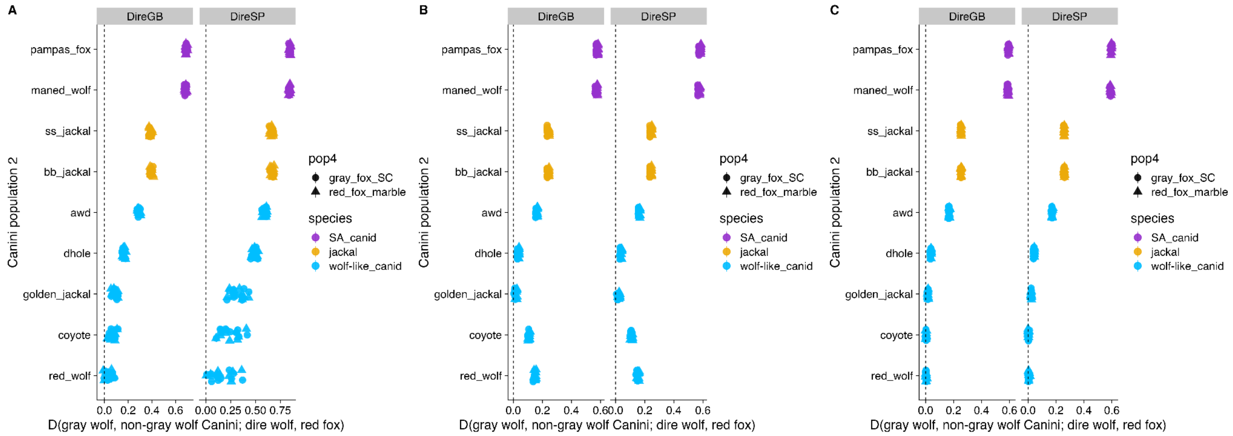


**Figure S19:** Comparison of *D*-statistics examining allele sharing between dire wolves and gray wolves calculated either using **A)** diploid genotypes called with *DeepVariant*, **B)** single-read pseudohaploid genotypes, or **C)** Consensify-style pseudohaploid genotypes. Diploid genotypes exhibit reference bias (excess allele sharing with gray wolves) which is exacerbated in the lower coverage SP individual, while single-read pseudohaploid genotypes have apparent outgroup attraction for taxa with higher sequencing error (coyote and red wolf, **Fig. S10**). The Consensify-style pseudohaploid genotypes appear to mitigate both sources of bias.
